# ProLoc: Text-guided Localization of Protein Functional Regions

**DOI:** 10.64898/2026.06.24.733131

**Authors:** Peishuo Liu, Jiaxin Fan, Mianzhi Pan, Jianbing Zhang

## Abstract

**Motivation:** Protein function is often mediated by specific sequence regions, such as domains, motifs and functional sites. Identifying these regions is important for understanding protein mechanisms, annotating newly sequenced proteins and prioritizing residues for experimental validation. However, existing protein function prediction and protein–text models mainly capture global protein-level associations, making it difficult to determine which residues support a given textual functional description. This limits their use for mechanistic interpretation and residue-level experimental prioritization.

**Results:** We introduce text-guided protein functional region localization, a span-level grounding task that identifies residue regions corresponding to natural-language functional descriptions. We construct an InterPro-derived localization benchmark of explicit protein–text–region examples, covering both domain-level and functional-site annotations with sequence-similarity-aware splits and a unified span-level evaluation protocol. We further propose ProLoc, a text-conditioned localization model built on raw ESM2-650M and PubMedBERT with direct residue-level localization and anchor-free span proposal generation. On the held-out test set, ProLoc substantially outperforms window-based adaptations of representative protein and protein–text models. Its direct output achieves the strongest single-region localization performance, reaching 0.7730 IoU@1, while its anchor-free proposal output improves visible multi-site recovery, reaching 0.9671 VM R@10 IoU50 and 0.9489 VM All-Hit@50.

**Availability and Implementation:** Source code and evaluation scripts are available at https://github.com/ShiDeng7rz/Proloc. The processed benchmark and data splits are archived at Zenodo: https://doi.org/10.5281/zenodo.20729714.

## Introduction

Protein function is often mediated by localized regions within a sequence, including domains, motifs and functional sites (Blum et al., 2021; Sigrist et al., 2026). Identifying such regions is essential for understanding protein mechanisms, annotating newly sequenced proteins and prioritizing residues for experimental validation. However, current protein function prediction benchmarks and sequence-based function predictors typically evaluate protein-level functional label assignment (Radivojac et al., 2013; Zhou et al., 2019; Kulmanov and Hoehndorf, 2020), while existing protein–text models are mainly optimized for global sequence–text association rather than residue-level grounding (Xu et al., 2023; Zhou et al., 2025; Su et al., 2024). This distinction is important: two proteins may share the same functional label while placing the corresponding residues at different positions, and short functional sites can be difficult to recover from a global protein-level prediction alone.

Recent protein language models have produced powerful sequence representations and have advanced a broad range of protein prediction tasks (Elnaggar et al., 2021; Lin et al., 2023). Meanwhile, protein–text models have started to align protein sequences with natural-language functional descriptions, enabling protein–text retrieval and semantic representation learning (Xu et al., 2023; Zhou et al., 2025; Su et al., 2025). Despite these advances, protein–text models are mainly optimized for global sequence–text association or retrieval-style matching, while protein language models generally provide sequence representations rather than text-conditioned residue grounding. As a result, they do not provide a direct formulation or evaluation protocol for localizing a functional text description to residue regions inside a given protein.

This gap motivates the task of text-guided protein functional region localization. Given a protein sequence and a natural-language functional description, the goal is to identify one or more residue spans corresponding to the described function. This task differs from standard protein function prediction because it requires locating the residues associated with a given functional description, rather than only assigning a protein-level functional label (Radivojac et al., 2013; Zhou et al., 2019). It also differs from protein–text retrieval, which ranks candidate proteins, peptides or pre-defined sequence fragments from a database, rather than predicting residue coordinates within a specified protein (Xu et al., 2023; Zhou et al., 2025; Su et al., 2025). Therefore, a unified benchmark is needed to evaluate whether protein and protein–text models can move beyond global semantic matching toward fine-grained functional grounding.

To this end, we construct an InterPro-derived benchmark for text-guided protein functional region localization using InterPro annotations and the corresponding UniProtKB protein sequences (Blum et al., 2021; UniProt Consortium, 2025). The benchmark contains both domain-level and functional-site annotations, paired with functional text descriptions and residue coordinates. We adopt a sequence-similarity-aware data split to reduce homology leakage, and evaluate all methods under a unified span-level protocol. Since existing protein and protein–text models are not directly designed to output residue spans for this task, we adapt representative models into a window-based grounding framework, where candidate sequence windows are ranked according to their similarity to the functional text query (Lin et al., 2023; Xu et al., 2023; Zhou et al., 2025; Su et al., 2025). This provides a consistent external comparison for retrieval-style grounding.

We further propose ProLoc, a text-conditioned residue localization model based on raw ESM2-650M (Lin et al., 2023) and PubMedBERT (Gu et al., 2021). ProLoc uses a shared text-guided residue fusion backbone that combines residue-level protein representations with both pooled and token-level text information. On top of this shared representation, we design two complementary output paths. The direct head predicts dense residue-level in-span scores and serves as the primary output for precise top-ranked region localization. The anchor-free proposal head (Tian et al., 2019) predicts start, end and length signals, which are decoded into ranked top-*K* span proposals. This design separates precise dense localization from structured proposal generation: the direct head provides accurate top-ranked localization, while the anchor-free head produces multiple candidate spans for top-*K* recovery.

Experiments show that ProLoc-Direct substantially improves over window-based adaptations of existing models. ProLoc-Direct achieves the strongest single-region localization performance on both domain and functional-site annotations. ProLoc-AF complements this by improving visible multi-site recovery, especially when the same functional description corresponds to multiple separated regions in the same protein. Importantly, ProLoc is trained without explicit multi-site group supervision; multi-site recovery is evaluated only at test time by grouping records with the same protein identifier and InterPro identifier.

In summary, our contributions are threefold. First, we define text-guided protein functional region localization as a span-level residue grounding task, and construct a large-scale InterPro-derived benchmark covering both domain and functional-site annotations. Second, we provide a unified comparison protocol that adapts representative protein representation models and protein–text models into window-based grounding baselines. Third, we propose ProLoc, a dual-output localization model that combines dense residue-level prediction with anchor-free span proposal generation, improving precise single-region localization while enabling structured top-*K* recovery of visible multi-site functional regions.

## Methods

### Task definition: text-guided protein functional region localization

We define text-guided protein functional region localization as identifying residue regions in a protein sequence conditioned on a functional text description. Given a protein sequence *P* = (*a*_1_, *a*_2_, …, *a*_*L*_) and a text query *q* describing a domain, motif or functional site, the model predicts one or more residue spans in *P* that support the queried function.

Each annotated region is represented as a half-open interval [*s, e*), where *s* is the start position and *e* is the exclusive end position. We use this span convention throughout training and evaluation. The span is also converted into a residue-level binary mask *y* ∈ {0, 1}^*L*^, where residues inside the annotated region are labeled as positives. Domain annotations usually correspond to longer continuous regions, whereas functional-site annotations are shorter and more sensitive to boundary shifts.

Unlike protein-level function prediction or protein–text retrieval, this task fixes the protein sequence and requires the model to localize which residue spans inside the protein support the given query. We evaluate two complementary settings. In single-region localization, each protein–text record has a target span, and the top-ranked prediction is evaluated against this span. In top-*K* proposal evaluation, the model returns a ranked list of candidate spans.

For visible multi-site evaluation, we group held-out test records sharing the same protein identifier and InterPro identifier, and evaluate whether ranked proposals recover multiple retained regions for the same protein–function pair. These groups are used only during evaluation. During training, each protein–text–region record is treated as an independent supervised example, without explicit multi-site group labels or set-level supervision.

### Construction of an InterPro-derived localization benchmark

We construct an InterPro-derived benchmark for text-guided protein functional region localization using InterPro annotations (Blum et al., 2021). Each raw record contains a protein sequence, an InterPro identifier, an entry type, a functional text description and residue-level coordinates. We retain annotations with valid coordinates and informative descriptions, and remove weakly specified entries containing terms such as “unknown”, “uncharacterized”, “putative” or DUF-like labels (Mistry et al., 2021) (Supplementary Methods S1.1). Five InterPro entry types are considered: `Domain`, `Active_site`, `Binding_site`, `Conserved_site` and `PTM`. Domain entries are treated as domain-level regions, whereas the remaining four types are grouped as functional-site regions.

The raw annotation pool contains 219.4 million domain records and 39.9 million site records (Supplementary Methods S1.1). We apply sequence- and region-level filtering to remove unsuitable localization targets. Protein lengths are restricted to 50–2000 amino acids. Domain regions are required to contain at least 20 residues, and regions covering more than 80% of the full protein are removed because they provide limited information for distinguishing localized functional regions from the full protein. Functional-site annotations follow the same protein-length constraint but do not use a minimum site-length threshold, since functional sites are often short. All final training and evaluation targets use the original InterPro residue coordinates.

To reduce the dominance of highly frequent InterPro identifiers, we use description-group-aware identifier sampling rather than raw-frequency sampling (Supplementary Fig. S1). Functional descriptions are normalized and grouped by recurrent description patterns, with high-frequency functional families manually consolidated and site annotations further stratified by subtype (Supplementary Methods S1.2). This procedure selects 1,200 domain identifiers and 450 functional-site identifiers. The selected identifiers and retained annotations are then converted into explicit protein–text–region examples. Duplicated annotation hits and near-identical regions within the same protein–InterPro pair are merged, followed by a per-pair retention rule: each protein–InterPro pair retains at most one domain span and up to two functional-site spans (Supplementary Methods S1.4). This yields 474,580 explicit positive examples, including 344,445 domain examples and 130,135 site examples.

To reduce homology leakage, we split the benchmark at the protein-cluster level. All candidate proteins are clustered using MMseqs2 at 50% sequence identity (Steinegger and Söding, 2017), and all proteins in the same cluster are assigned to the same split. The split is constructed to balance protein counts, sample counts, the domain/site ratio and the functional-site subtype distribution. We also audit InterPro identifier support across splits to avoid evaluation subsets dominated by poorly represented identifiers (Supplementary Methods S1.5). After cluster-level assignment and split auditing, the final benchmark contains 407,986 protein–text–region examples, including 353,555 training examples, 4,999 validation examples and 49,432 held-out test examples. The validation subset is used for checkpoint selection, and all final comparisons are reported on the held-out test set. Table 1 summarizes the final statistics.

**Table 1.** Statistics of the InterPro-derived localization benchmark.

| Split | Samples | Proteins | Domain | Site | InterPro IDs |
| --- | --- | --- | --- | --- | --- |
| Train | 353,555 | 268,334 | 256,727 | 96,828 | 1,650 |
| Valid-5k | 4,999 | 4,804 | 3,717 | 1,282 | 1,477 |
| Test | 49,432 | 34,219 | 36,547 | 12,885 | 1,477 |

During training, each protein–text–region example is treated independently, without explicit multi-site group labels or set-level supervision. For evaluation, we additionally construct visible multisite groups by grouping test examples with the same protein identifier and InterPro identifier (Supplementary Methods S1.6 and S4.4). The held-out test set contains 1,731 visible multi-site groups, all from functional-site annotations (Supplementary Table S2). These groups are used only to evaluate whether ranked span proposals can recover multiple retained functional regions for the same protein–function pair.

### Baseline adaptation and unified comparison protocol

Existing protein representation models and protein–text models are not originally designed to produce text-conditioned residue-level localization maps. We therefore adapt representative models to our benchmark through a unified region–text alignment and window-based grounding protocol. The evaluated baselines include Raw ESM2-650M (Lin et al., 2023), ProtBERT and ProtT5 (Elnaggar et al., 2021), ProtST (Xu et al., 2023), ProtCLIP (Zhou et al., 2025) and ProTrek (Su et al., 2025), covering protein-only language models, protein–text pretrained models and sequence–structure– text multimodal representations. All baselines use the same data splits, validation-based checkpoint selection and span-level evaluator (Supplementary Methods S2).

During adaptation, each annotated protein–text–region example is converted into a positive region–text pair. The annotated residue span is encoded as a protein region by attention pooling over residue features, and the corresponding InterPro description is used as the text query. Protein-only encoders are paired with PubMedBERT as the text tower, whereas protein–text pretrained models use their available text encoders when applicable. We train lightweight adaptation modules with a CLIP-style in-batch contrastive objective (Radford et al., 2021; Chen et al., 2020), where matched region–text pairs are positives and mismatched pairs in the same mini-batch are negatives. Examples with the same normalized functional description are masked out from the negative set to avoid false-negative penalties (Chuang et al., 2020). The pretrained encoders are kept frozen, and only projection heads, region attention pooling and temperature parameters are optimized.

At inference time, each adapted baseline is evaluated using window-based grounding (Supplementary Methods S2.3). Given a full-length protein sequence and a functional text query, candidate sequence windows are generated along the protein sequence, scored by region– text similarity and ranked as predicted spans. The top-ranked window is used for IoU@1 and boundary-based metrics, while the top-10 windows are used for Best@10 and visible multi-site evaluation (Supplementary Methods S4.4). In contrast, ProLoc directly predicts dense residue-level localization scores and anchor-free span proposals. For fair comparison, all methods are converted into the same ranked span format and evaluated with the same span-level metrics, so that the comparison is not confounded by heterogeneous output formats or evaluation procedures.

### Proposed model architecture

We propose ProLoc, a text-conditioned residue localization model that maps a protein sequence and a functional text query to both dense residue-level localization scores and ranked span proposals. The model consists of a protein encoder, a text encoder, a shared text-guided residue fusion backbone and two complementary output heads. The direct localization head is designed for precise single-region grounding, whereas the anchor-free proposal head provides a structured top-*K* span proposal interface. An overview of the architecture is shown in Fig. 1.

**Figure 1.**
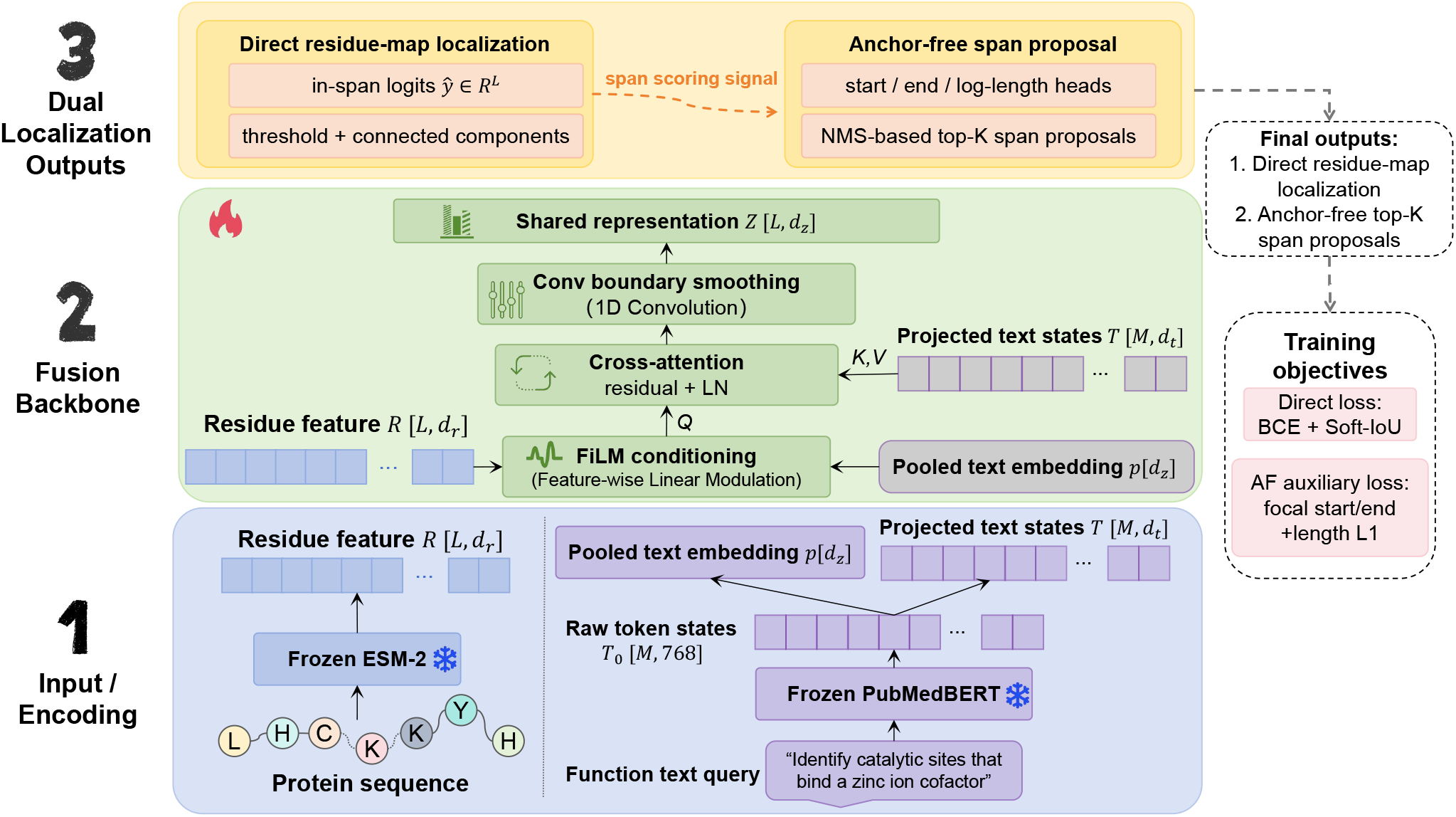
Overview of ProLoc. Given a protein sequence and a functional text query, ProLoc encodes residue-level protein representations using raw ESM2-650M and text representations with PubMedBERT. A shared text-guided residue fusion backbone integrates pooled text semantics and token-level text evidence through feature-wise linear modulation (FiLM), residue-to-text cross-attention and a one-dimensional convolutional refinement module. The direct localization head predicts dense residue-level in-span scores for precise single-region localization, while the anchor-free proposal head predicts start, end and length signals to generate ranked top-*K* span proposals for structured top-*K* recovery.

Given a protein sequence *P* = (*a*_1_, *a*_2_, …, *a*_*L*_), where *L* is the sequence length, the protein encoder outputs residue-level representations

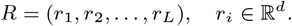

In our main model, we use raw ESM2-650M as the protein encoder (Lin et al., 2023). Given a functional text query *q*, the text encoder outputs a pooled text embedding *t*_*p*_, which summarizes the global query semantics, and token-level text states

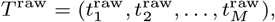

where *M* is the number of text tokens. We use PubMedBERT as the text encoder (Gu et al., 2021). Since the dimensionality of the text states may differ from that of the residue features, the token-level text states are projected into the residue representation space:

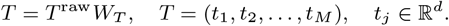

The shared residue fusion backbone incorporates both global and token-level text information. First, we apply feature-wise linear modulation (FiLM) to condition each residue representation on the pooled text embedding (Perez et al., 2018):

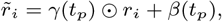

where *γ*(·) and *β*(·) are text-conditioned affine transformations and ⊙ denotes element-wise multiplication. This step injects global functional semantics into residue-level protein features, enabling the model to adapt residue representations according to the queried function.

To capture finer-grained residue–text interactions, we then apply residue-to-text cross-attention (Vaswani et al., 2017). The FiLM-modulated residue features serve as queries, and the projected token-level text states serve as keys and values:

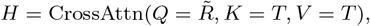

where 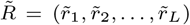 and *H* ∈ ℝ^*L*×*d*^. The attention output is added back to the FiLM-modulated residue features through a residual connection and layer normalization, allowing token-level query information to be incorporated differently across residues.

The fused residue features are further processed by a lightweight one-dimensional convolutional tower:

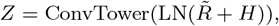

where *Z* = (*z*_1_, *z*_2_, …, *z*_*L*_) denotes the final text-conditioned residue representation. The convolutional tower aggregates local sequence context and smooths boundary-level evidence, which is useful for span-level localization.

On top of *Z*, the direct localization head predicts a dense in-span logit for each residue:

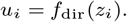

The resulting residue-level score map is decoded by thresholding and connected-component extraction (Supplementary Methods S4.2). This direct output serves as the primary prediction for single-region localization, where the goal is to identify the most relevant functional span for a given protein–text pair.

In parallel, the anchor-free proposal head predicts three residue-level signals:

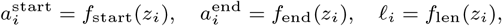

where 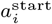 and 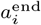 are start and end logits for residue *i*, and *ℓ*_*i*_ predicts the log-length of a span starting at residue *i*. During decoding, start and end scores provide boundary evidence, length prediction provides a span-size prior, and the dense direct scores provide span-level confidence (Supplementary Methods S4.3).

Candidate spans are generated from the boundary and length signals, ranked by their combined confidence, and filtered by non-maximum suppression to produce top-*K* proposals.

The two output paths are complementary. The direct head produces a dense residue probability map and serves as the primary output for accurate top-1 localization. The anchor-free head explicitly models boundary and length information, making it better suited for structured top-*K* proposal generation. Both heads share the same text-guided residue representation, allowing ProLoc to jointly learn residue-level localization and span-level proposal signals.

### Training objectives

ProLoc is trained with residue-level localization supervision and auxiliary anchor-free span supervision. For each protein–text–region example, the annotated region is represented as a half-open interval [*s, e*), where *s* is the start position and *e* is the exclusive end position.

The span is converted into a residue-level binary mask *y* ∈ {0, 1}^*L*^, where residues inside the annotated region are treated as positives, other valid residues are treated as annotation-relative negatives for the retained training example, and padding positions are excluded from loss computation (Supplementary Methods S3.3).

The direct localization head predicts dense in-span logits *u* = (*u*_1_, …, *u*_*L*_). Let *p* = *σ*(*u*) denote the residue-level in-span probabilities. We supervise this head with a combination of masked residue-level binary cross-entropy and soft-IoU loss (Rahman and Wang, 2016):

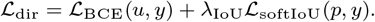

The BCE term encourages accurate residue-wise classification, whereas the soft-IoU term provides a differentiable overlap-oriented objective between the predicted residue map and the annotated span. This direct loss is the primary localization objective.

For the anchor-free proposal branch, we construct soft boundary targets for the start and end heads. The start target is a Gaussian-shaped soft label centered at *s*, and the end target is centered at *e* − 1, corresponding to the last residue in the half-open interval [*s, e*). The start and end heads are trained with focal binary cross-entropy losses (Lin et al., 2017):

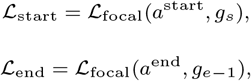

where *g*_*s*_ and *g*_*e*−1_ denote the soft boundary targets for the start and end positions, respectively. Soft boundary supervision reduces over-penalizing near-boundary predictions and provides smoother training signals, which is especially useful for short functional sites.

The length head is supervised only at the ground-truth start position, because the predicted span length has a clear interpretation only when conditioned on a valid start. The target log-length is defined as

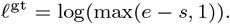

The length loss is

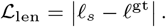

This term provides a span-size prior for anchor-free proposal decoding.

The final training objective combines the direct localization loss and the anchor-free auxiliary losses:

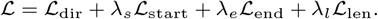

Here, *λ*_*s*_, *λ*_*e*_, and *λ*_*l*_ control the contributions of the anchor-free auxiliary losses (Supplementary Table S5). The direct loss remains the main localization objective, while the anchor-free losses provide additional boundary and span-size supervision to the shared text-guided residue representation.

During training, each protein–text–region example is treated independently. No explicit multi-site group label or set-level supervision is used. Thus, the model learns residue-level and span-level proposal signals from individual annotated regions, while multisite recovery is evaluated only at test time through ranked top-*K* span proposal metrics.

### Evaluation metrics

All methods are evaluated under a unified span-level protocol (Supplementary Methods S4.1). For each protein–text pair, each method output is normalized into a ranked list of predicted spans {*Ŝ*_1_, *Ŝ*_2_, …, *Ŝ*_*K*_}. For window-based baselines, ranked spans are candidate windows; for ProLoc-Direct, spans are decoded from dense residue-level in-span scores; and for ProLoc-AF, spans are decoded from anchor-free start, end, and length predictions. All decoded spans are compared with the same evaluator.

For single-region localization, we use the intersection-over-union between the top-ranked prediction *Ŝ*_1_ and the ground-truth span *S*:

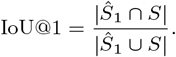

We report IoU@1 on the full test set and separately on domain and functional-site annotations. Since short functional-site regions are sensitive to small boundary shifts, we also report boundary mean absolute error:

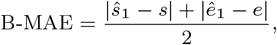

where (*ŝ*_1_, *ê*_1_) and (*s, e*) denote the predicted and ground-truth span boundaries, respectively. Lower B-MAE indicates more accurate boundary localization.

To evaluate candidate-set quality beyond the top-1 prediction, we report Best@10:

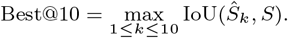

This metric reflects the quality of the ranked proposal set and is useful for candidate-set evaluation beyond the top-1 prediction. We also report Best@10 separately for domain and functional-site annotations (Supplementary Methods S4.4).

For visible multi-site evaluation, we group test records with the same protein identifier and InterPro identifier. Each group contains the benchmark-visible ground-truth regions retained for the same protein–function pair. Given the top-10 predicted spans, VM R@10 IoU50 measures the fraction of visible ground-truth regions that are matched by at least one prediction with IoU ≥ 0.5. VM R@10 IoU70 is defined similarly with an IoU threshold of 0.7. VM All-Hit@50 is a group-level metric: a group is counted as successful only if every visible ground-truth region in that group is matched by at least one top-10 prediction with IoU ≥ 0.5. VM Mean IoU@10 averages, over all visible ground-truth regions, the best IoU achieved by any top-10 prediction.

These metrics capture complementary aspects of the task. IoU@1 and B-MAE evaluate precise single-region localization, Best@10 evaluates the quality of the ranked candidate set, and visible multi-site metrics evaluate whether a model can recover multiple benchmark-visible functional regions for the same protein– function query. Residue-level diagnostic metrics and decoding hyperparameters are reported in the Supplementary Methods S4.

### Implementation details

All models are implemented in PyTorch (Paszke et al., 2019). For ProLoc, raw ESM2-650M and PubMedBERT are kept frozen, and only the text-guided fusion backbone and localization heads are optimized. Window-based baselines also keep pretrained encoders frozen and train only lightweight region–text adaptation modules. Checkpoints are selected on the validation subset, and all final results are reported on the held-out test set using the same span-level evaluator. Decoding settings, window-generation parameters, non-maximum suppression parameters and loss hyperparameters are reported in Supplementary Methods S2–S4.

## Results

### Window-based baselines remain limited in top-1 localization

We first evaluate how existing protein representation and protein– text models perform on text-guided functional region localization when adapted through window-based grounding. For each adapted baseline, candidate sequence windows are ranked according to their similarity to the functional text query, and the ranked windows are compared with the annotated residue span. This setting provides a natural evaluation protocol for models without native text-conditioned residue-level localization outputs.

As shown in Table 2, window-based baselines achieve moderate top-10 candidate-set performance but remain limited in top-1 localization accuracy. Among the external baselines, ProtT5 and ProtCLIP achieve the highest IoU@1 scores, reaching 0.5498 and 0.5439, respectively. Their Best@10 scores increase to 0.7626 and 0.7606, indicating that the top-ranked candidate set often contains windows with substantially better overlap than the top-1 prediction. However, the gap between IoU@1 and Best@10 also suggests that window-based ranking does not reliably place the best-overlapping region first, which limits its usefulness for precise functional grounding.

**Table 2.** Main results on the held-out test set. Window-based baselines rank candidate sequence windows according to region–text similarity. ProLoc-Direct decodes spans from dense residue-level in-span scores, whereas ProLoc-AF decodes ranked proposals from anchor-free start, end and length signals, with dense direct scores used for span-level confidence. Higher is better except for B-MAE.

| Method | Type | IoU@1 | Domain IoU@1 | Site IoU@1 | B-MAE ↓ | Best@10 | Domain Best@10 | Site Best@10 | VM R@10 IoU50 | VM R@10 IoU70 | VM All-Hit@50 | VM Mean IoU@10 |
| --- | --- | --- | --- | --- | --- | --- | --- | --- | --- | --- | --- | --- |
| Raw ESM2 | Window | 0.4544 | 0.4483 | 0.4720 | 116.72 | 0.7164 | 0.7245 | 0.6935 | 0.8304 | 0.5859 | 0.7063 | 0.6638 |
| ProtST | Window | 0.4870 | 0.4857 | 0.4908 | 113.06 | 0.7332 | 0.7466 | 0.6951 | 0.8285 | 0.5970 | 0.7149 | 0.6603 |
| ProtBERT | Window | 0.4806 | 0.4883 | 0.4590 | 117.61 | 0.7283 | 0.7473 | 0.6747 | 0.7980 | 0.5889 | 0.6519 | 0.6396 |
| ProtT5 | Window | 0.5498 | 0.5716 | 0.4876 | 101.90 | 0.7626 | 0.7845 | 0.7003 | 0.8444 | 0.5975 | 0.7405 | 0.6738 |
| ProtCLIP | Window | 0.5439 | 0.5615 | 0.4938 | 104.72 | 0.7606 | 0.7821 | 0.6996 | 0.8481 | 0.6239 | 0.7343 | 0.6780 |
| ProTrek | Window | 0.5213 | 0.5314 | 0.4926 | 105.88 | 0.7449 | 0.7646 | 0.6890 | 0.8177 | 0.5683 | 0.7106 | 0.6495 |
| Direct-only ProLoc | Direct | 0.7621 | 0.7798 | 0.7120 | 68.51 | 0.8061 | 0.8025 | 0.8113 | 0.8171 | 0.7466 | 0.7408 | 0.7524 |
| ProLoc-Direct | Direct | **0.7730** | **0.7938** | 0.7141 | **65.77** | 0.8142 | **0.8153** | 0.8163 | 0.8322 | 0.7602 | 0.7438 | 0.7597 |
| ProLoc-AF | AF proposal | 0.7348 | 0.7511 | 0.6886 | 74.79 | 0.8062 | 0.7997 | **0.8246** | **0.9671** | **0.8741** | **0.9489** | **0.8276** |

The limitation is more pronounced for short functional-site regions. For the stronger window-based baselines, performance is higher on domain annotations than on functional-site annotations, reflecting the difficulty of localizing compact regions with fixed multi-scale candidate windows. For example, ProtT5 achieves 0.5716 Domain IoU@1 compared with 0.4876 Site IoU@1. These results suggest that although pretrained protein and protein–text representations provide useful semantic signals, window-based grounding is insufficient as a stand-alone strategy for accurate top-1 residue-level localization, motivating direct text-conditioned localization in ProLoc.

### Direct localization improves single-region grounding

We next evaluate whether directly predicting residue-level localization maps improves text-guided functional region grounding. Unlike window-based baselines, the direct head predicts dense in-span scores over protein residues and converts high-scoring connected components into span predictions. We denote the separately trained model with only the direct localization head as Direct-only ProLoc. In the full model, ProLoc-Direct denotes the direct-head output decoded from dense residue-level in-span logits after joint training with anchor-free auxiliary objectives.

As shown in Table 2, direct residue-level localization substantially improves top-1 single-region grounding over window-based baselines. Direct-only ProLoc achieves 0.7621 IoU@1, whereas the strongest external baseline, ProtT5, reaches 0.5498 IoU@1. This improvement is consistent across annotation types: Direct-only ProLoc obtains 0.7798 Domain IoU@1 and 0.7120 Site IoU@1, compared with 0.5716 and 0.4876 for ProtT5. These results indicate that text-conditioned residue-level prediction is more effective for top-1 fine-grained functional grounding than window-based ranking.

As shown in Table 3, joint training with anchor-free auxiliary objectives further improves the direct output over Direct-only ProLoc. Compared with Direct-only ProLoc, ProLoc-Direct increases IoU@1 from 0.7621 to 0.7730 and reduces B-MAE from 68.51 to 65.77, indicating more accurate top-ranked spans and improved boundary localization. The strongest performance is observed for domain localization, where ProLoc-Direct reaches 0.7938 Domain IoU@1 and 0.8153 Domain Best@10. These results suggest that dense residue-level supervision provides a strong localization signal, while auxiliary span-level training further regularizes the shared text-conditioned residue representation.

**Table 3.** Ablation and control analysis of supervised dense localization, anchor-free auxiliary training and the length branch. Higher is better except for B-MAE.

| Variant | Output | IoU@1 | B-MAE ↓ | Best@10 | VM R@10 IoU50 | VM R@10 IoU70 | VM All-Hit@50 | VM Mean IoU@10 |
| --- | --- | --- | --- | --- | --- | --- | --- | --- |
| Raw-FiLM | Direct | 0.5568 | 99.59 | 0.6023 | 0.3645 | 0.1521 | 0.1810 | 0.3923 |
| Direct-only ProLoc | Direct | 0.7621 | 68.51 | 0.8061 | 0.8171 | 0.7466 | 0.7408 | 0.7524 |
| ProLoc-Direct | Direct | **0.7730** | **65.77** | 0.8142 | 0.8322 | 0.7602 | 0.7438 | 0.7597 |
| ProLoc-AF | AF proposal | 0.7348 | 74.79 | 0.8062 | **0.9671** | **0.8741** | **0.9489** | **0.8276** |
| ProLoc-AF w/o length | Direct | 0.7655 | 69.50 | 0.8074 | 0.8168 | 0.7522 | 0.7112 | 0.7519 |
| ProLoc-AF w/o length | AF proposal | 0.7702 | 69.68 | **0.8327** | 0.8119 | 0.7842 | 0.6983 | 0.7456 |

It is worth noting that the direct head remains the strongest output for precise single-region localization. Compared with the anchor-free proposal output, ProLoc-Direct achieves higher IoU@1 and lower B-MAE. We therefore use the direct head as the primary output for single-region localization and analyze the anchor-free head separately as a structured top-*K* proposal mechanism in the next section.

### Anchor-free proposals improve visible multi-site recovery

Although the direct head provides the strongest single-region localization, some functional-site annotations are not always represented by a single continuous region. In our benchmark, visible multi-site groups arise from site annotations, where the same functional description may correspond to multiple short and separated regions within one protein. Decoding such cases from a dense residue score map relies on thresholding and connected-component extraction, which works well for top-1 localization but is not explicitly optimized for ranked multi-span proposal generation.

ProLoc-AF is designed to address this limitation by decoding spans from start, end and length predictions. This output is complementary to ProLoc-Direct: the direct head focuses on precise single-region localization, whereas the anchor-free head provides structured top-*K* span proposals. As shown in Table 2, ProLoc-AF achieves lower overall IoU@1 than ProLoc-Direct, with 0.7348 compared with 0.7730. However, it achieves the best Site Best@10 score of 0.8246, indicating that the anchor-free output provides stronger ranked candidate sets for short functional-site regions.

The advantage of ProLoc-AF is most evident in visible multisite recovery. Compared with ProLoc-Direct, ProLoc-AF increases VM R@10 IoU50 from 0.8322 to 0.9671 and VM R@10 IoU70 from 0.7602 to 0.8741. It also improves VM All-Hit@50 from 0.7438 to 0.9489 and VM Mean IoU@10 from 0.7597 to 0.8276. These results indicate that anchor-free decoding provides a more suitable interface for recovering multiple visible functional-site regions associated with the same protein–function query. Therefore, ProLoc-Direct and ProLoc-AF serve different purposes: the former serves precise top-1 localization, while the latter is used for structured top-*K* proposals for visible multi-region recovery.

Representative qualitative examples and failure cases are provided in Supplementary Figs. S3 and S4, respectively, further illustrating the complementary behavior of the direct and anchor-free outputs.

### Ablation and control analysis

We further analyze the roles of supervised dense localization, anchor-free auxiliary objectives and the length branch. We first include Raw-FiLM as a supervised dense control that uses the same raw ESM2-650M and PubMedBERT encoders, but conditions residue representations only through pooled-text FiLM modulation, without token-level residue-to-text cross-attention or anchor-free proposal learning. As shown in Table 3, Raw-FiLM reaches 0.5568 IoU@1 and 0.6023 Best@10, clearly below Direct-only ProLoc and ProLoc-Direct. This indicates that the gain of ProLoc is not explained solely by replacing window ranking with dense residue-level supervision.

We next compare the two outputs decoded from the same full ProLoc checkpoint. ProLoc-Direct is stronger for precise single-region localization, whereas ProLoc-AF is substantially better for visible multi-site recovery. This indicates that the two heads serve different roles rather than competing as alternative decoders: the direct head provides the most accurate top-1 localization, while the anchor-free head provides a structured ranked proposal set for recovering multiple visible functional regions.

Finally, we examine the effect of the length branch. Removing length supervision still yields strong top-1 and Best@10 scores for the AF output, with 0.7702 IoU@1 and 0.8327 Best@10. However, its multi-site recovery drops sharply: VM R@10 IoU50 decreases from 0.9671 to 0.8119, and VM All-Hit@50 decreases from 0.9489 to 0.6983. These results suggest that the length branch mainly provides a span-size prior that stabilizes top-*K* proposal generation, especially when multiple functional sites must be recovered within the same protein–function query.

A matched wrong-query control further confirms query dependence: replacing each test query with a type- and length-matched incorrect functional description reduces ProLoc-Direct from 0.7730 to 0.1621 IoU@1 and from 0.8142 to 0.1930 Best@10, indicating that the model does not simply highlight fixed protein regions. Full control metrics are reported in Supplementary Tables S7 and S8.

## Discussion

We introduced text-guided protein functional region localization as a span-level grounding task that links functional text descriptions to residue regions within a given protein sequence. The InterPro-derived benchmark and unified span-level evaluation protocol provide a controlled setting for evaluating this fine-grained protein–text grounding ability across domain and functional-site annotations.

Our results show that window-based adaptations of existing protein representation and protein–text models provide useful window-based grounding baselines, but remain limited for precise top-1 localization. In contrast, direct residue-level localization substantially improves single-region grounding, indicating that dense text-conditioned residue prediction is beneficial for this task. The two ProLoc outputs serve complementary roles: the direct head achieves the strongest single-region localization and boundary accuracy, whereas the anchor-free proposal head provides a more effective top-*K* interface for visible multi-site recovery. The ablation results further suggest that anchor-free supervision benefits the shared residue representation, while the length branch mainly stabilizes multi-span proposal generation.

Several limitations remain. The benchmark is derived from InterPro (Blum et al., 2021) and therefore inherits its annotation coverage and biases; unannotated functional regions are not available as positive labels and should not be interpreted as confirmed negatives. The current benchmark assumes that the queried function is present in the protein; abstention or absent-query detection is outside the current scope. In addition, ProLoc is trained on independent protein–text–region examples without explicit multisite group supervision, although visible multi-site recovery is evaluated at test time. Future work could incorporate set-level objectives, structural context, or additional experimentally validated annotations to improve localization of discontinuous or spatially organized functional-site regions.

## Data availability

Source code and evaluation scripts are available at https://github.com/ShiDeng7rz/Proloc. The processed benchmark and data splits are archived at Zenodo: https://doi.org/10.5281/zenodo.20729714.

## References

M. Blum, H.-Y. Chang, S. Chuguransky, T. Grego, S. Kandasaamy, A. Mitchell, G. Nuka, T. Paysan-Lafosse, M. Qureshi, S. Raj, et al. The interpro protein families and domains database: 20 years on. Nucleic acids research, 49 (D1):D344–D354, 2021.

T. Chen, S. Kornblith, M. Norouzi, and G. Hinton. A simple framework for contrastive learning of visual representations. In International conference on machine learning, pages 1597–1607. PmLR, 2020.

C.-Y. Chuang, J. Robinson, Y.-C. Lin, A. Torralba, and S. Jegelka. Debiased contrastive learning. Advances in neural information processing systems, 33:8765–8775, 2020.

A. Elnaggar, M. Heinzinger, C. Dallago, G. Rehawi, Y. Wang, L. Jones, T. Gibbs, T. Feher, C. Angerer, M. Steinegger, et al. Prottrans: toward understanding the language of life through self-supervised learning. IEEE transactions on pattern analysis and machine intelligence, 44(10):7112–7127, 2021.

Y. Gu, R. Tinn, H. Cheng, M. Lucas, N. Usuyama, X. Liu, T. Naumann, J. Gao, and H. Poon. Domain-specific language model pretraining for biomedical natural language processing. ACM Transactions on Computing for Healthcare (HEALTH), 3(1):1–23, 2021.

M. Kulmanov and R. Hoehndorf. Deepgoplus: improved protein function prediction from sequence. Bioinformatics, 36(2):422–429, 2020.

T.-Y. Lin, P. Goyal, R. Girshick, K. He, and P. Dollár. Focal loss for dense object detection. In Proceedings of the IEEE international conference on computer vision, pages 2980–2988, 2017.

Z. Lin, H. Akin, R. Rao, B. Hie, Z. Zhu, W. Lu, N. Smetanin, R. Verkuil, O. Kabeli, Y. Shmueli, et al. Evolutionary-scale prediction of atomic-level protein structure with a language model. Science, 379(6637):1123–1130, 2023.

J. Mistry, S. Chuguransky, L. Williams, M. Qureshi, G. A. Salazar, E. L. Sonnhammer, S. C. Tosatto, L. Paladin, S. Raj, L. J. Richardson, et al. Pfam: The protein families database in 2021. Nucleic acids research, 49(D1):D412–D419, 2021.

A. Paszke, S. Gross, F. Massa, A. Lerer, J. Bradbury, G. Chanan, T. Killeen, Z. Lin, N. Gimelshein, L. Antiga, A. Desmaison, A. Köpf, E. Yang, Z. DeVito, M. Raison, A. Tejani, S. Chilamkurthy, B. Steiner, L. Fang, J. Bai, and S. Chintala. Pytorch: An imperative style, high-performance deep learning library. In Advances in Neural Information Processing Systems, volume 32, pages 8024–8035, 2019.

E. Perez, F. Strub, H. De Vries, V. Dumoulin, and A. Courville. Film: Visual reasoning with a general conditioning layer. In Proceedings of the AAAI conference on artificial intelligence, volume 32, 2018.

A. Radford, J. W. Kim, C. Hallacy, A. Ramesh, G. Goh, S. Agarwal, G. Sastry, A. Askell, P. Mishkin, J. Clark, et al. Learning transferable visual models from natural language supervision. In International conference on machine learning, pages 8748–8763. PmLR, 2021.

P. Radivojac, W. T. Clark, T. R. Oron, A. M. Schnoes, T. Wittkop, A. Sokolov, K. Graim, C. Funk, K. Verspoor, A. Ben-Hur, et al. A large-scale evaluation of computational protein function prediction. Nature methods, 10(3):221–227, 2013.

M. A. Rahman and Y. Wang. Optimizing intersection-over-union in deep neural networks for image segmentation. In International symposium on visual computing, pages 234–244. Springer, 2016.

C. J. Sigrist, B. A. Cuche, E. de Castro, E. Coudert, N. Redaschi, and A. Bridge. The prosite database for protein families, domains, and sites. Nucleic Acids Research, 54(D1):D451–D458, 2026.

M. Steinegger and J. Söding. Mmseqs2 enables sensitive protein sequence searching for the analysis of massive data sets. Nature biotechnology, 35(11):1026–1028, 2017.

J. Su, X. Zhou, X. Zhang, and F. Yuan. Protrek: Navigating the protein universe through tri-modal contrastive learning. bioRxiv, pages 2024–05, 2024.

J. Su, Y. He, S. You, S. Jiang, X. Zhou, X. Zhang, Y. Wang, X. Su, I. Tolstoy, X. Chang, et al. A trimodal protein language model enables advanced protein searches. Nature Biotechnology, pages 1–7, 2025.

Z. Tian, C. Shen, H. Chen, and T. He. Fcos: Fully convolutional one-stage object detection. In Proceedings of the IEEE/CVF international conference on computer vision, pages 9627–9636, 2019.

UniProt Consortium. Uniprot: the universal protein knowledgebase in 2025. Nucleic Acids Research, 53(D1): D609–D617, 2025. doi: 10.1093/nar/gkae1010.

A. Vaswani, N. Shazeer, N. Parmar, J. Uszkoreit, L. Jones, A. N. Gomez, L-. Kaiser, and I. Polosukhin. Attention is all you need. Advances in neural information processing systems, 30, 2017.

M. Xu, X. Yuan, S. Miret, and J. Tang. Protst: Multi-modality learning of protein sequences and biomedical texts. In International conference on machine learning, pages 38749–38767. PMLR, 2023.

H. Zhou, M. Yin, W. Wu, M. Li, K. Fu, J. Chen, J. Wu, and Z. Wang. Protclip: Function-informed protein multi-modal learning. In Proceedings of the AAAI Conference on Artificial Intelligence, volume 39, pages 22937–22945, 2025.

N. Zhou, Y. Jiang, T. R. Bergquist, A. J. Lee, B. Z. Kacsoh, A. W. Crocker, K. A. Lewis, G. Georghiou, H. N. Nguyen, M. N. Hamid, et al. The cafa challenge reports improved protein function prediction and new functional annotations for hundreds of genes through experimental screens. Genome biology, 20(1):244, 2019.

