## Supplementary Material for "ProLoc: Text-guided Localization of Protein Functional Regions"

#### Supplementary Methods

##### S1. Benchmark construction and split control

###### S1.1 Source annotations and filtering

We constructed the localization benchmark from InterPro-derived functional region annotations. We used InterPro release 108.0, released on 2026-01-29, together with the corresponding UniProtKB release `2026_01`, released on 2026-01-28. After linking InterPro annotations to UniProtKB protein sequences (Blum et al., 2021; UniProt Consortium, 2025), each retained record contained a full-length protein sequence, an InterPro identifier, an entry type, a functional text description and residue-level coordinates. Since the goal of this benchmark is to evaluate functional region localization rather than protein-level function classification, we retained only records with valid sequence coordinates and available full-length protein sequences.

We considered two broad categories of functional regions: protein domains and functional sites. Five InterPro entry types were retained: `Domain`, `Active_site`, `Binding_site`, `Conserved_site` and `PTM`. `Domain` entries were treated as domain-level regions, whereas the other four entry types were grouped as site-level regions. Before the subsequent description- and region-level filters, the retained entry-type annotation pool contained approximately 219.4 million domain annotation records and 39.9 million functional-site annotation records. Other InterPro entry types, such as family- or superfamily-level entries, were not included because they are less directly tied to a specific localized residue span under our benchmark definition.

To improve text-label quality, we removed records whose functional descriptions were weakly specified or uninformative. After lower-casing and normalizing functional descriptions, entries containing terms such as "unknown", "uncharacterized", "putative" or DUF-like labels (Mistry et al., 2021) were excluded. This step favors better-characterized annotations, but reduces textual supervision that does not provide a clear functional localization target.

We further applied sequence- and region-level filtering rules. Full-length protein sequences were restricted to 50–2,000 amino acids. For domain annotations, the annotated region was required to contain at least 20 residues, and regions covering more than 80% of the full protein sequence were removed, because such near-full-length regions provide limited information for distinguishing localized functional regions from the whole protein. Functional-site annotations used the same protein-length constraint but did not use a minimum site-length threshold, since many functional sites are inherently short.

Duplicated annotation hits and near-duplicate regions within the same protein–InterPro pair were merged using deterministic preprocessing rules. For two spans, we defined the overlap ratio as the intersection length divided by the shorter span length. Two functional-site spans were treated as near-duplicates if their overlap ratio was at least 0.80, or if $|\Delta s| \leq 5$ and $|\Delta e| \leq 5$. Two domain spans were treated as near-duplicates if their overlap ratio was at least 0.90, or if both their start and end coordinates differed by no more than 10 residues. We then constructed a near-duplicate graph within each protein–InterPro pair and collapsed each connected component into a single representative span, selected as the first span after deterministic sorting. This procedure collapsed 2,157 redundant annotation spans across 1,277 near-duplicate components. All final training and evaluation targets used the original InterPro residue coordinates of the retained representative spans. Therefore, the reported localization metrics are computed against the original InterPro annotation boundaries rather than artificially padded spans.

All residue coordinates were converted before model training and evaluation into a unified 0-based half-open span representation $[s, e)$, where $s$ denotes the start residue position and $e$ denotes the exclusive end position. This coordinate convention was used to construct residue-level masks, decode predicted spans and compute all span-level overlap metrics.

##### S1.2 Description-based identifier sampling

After source annotation filtering, we sampled functional identifiers for benchmark construction. InterPro annotations are highly long-tailed: a small number of identifiers occur in many proteins, whereas many identifiers have limited support. Directly sampling identifiers according to raw annotation frequency would make the benchmark dominated by a few abundant domain families or common functional descriptions. Our goal was therefore not to reproduce the natural InterPro frequency distribution, but to construct a localization benchmark with broader functional coverage, more balanced domain–site composition and improved representation of both frequent and less frequent functional groups. This imbalance was confirmed at the level of normalized-description functional groups (Supplementary Fig. S1). In the raw annotation pool, the top 1% of domain groups accounted for 44.1% of domain annotations, and the top 10% accounted for 77.5%. Site annotations were also long-tailed, with the top 10% of functional-site groups covering 59.8% of site annotations. These statistics motivated the coverage-oriented sampling strategy described below.

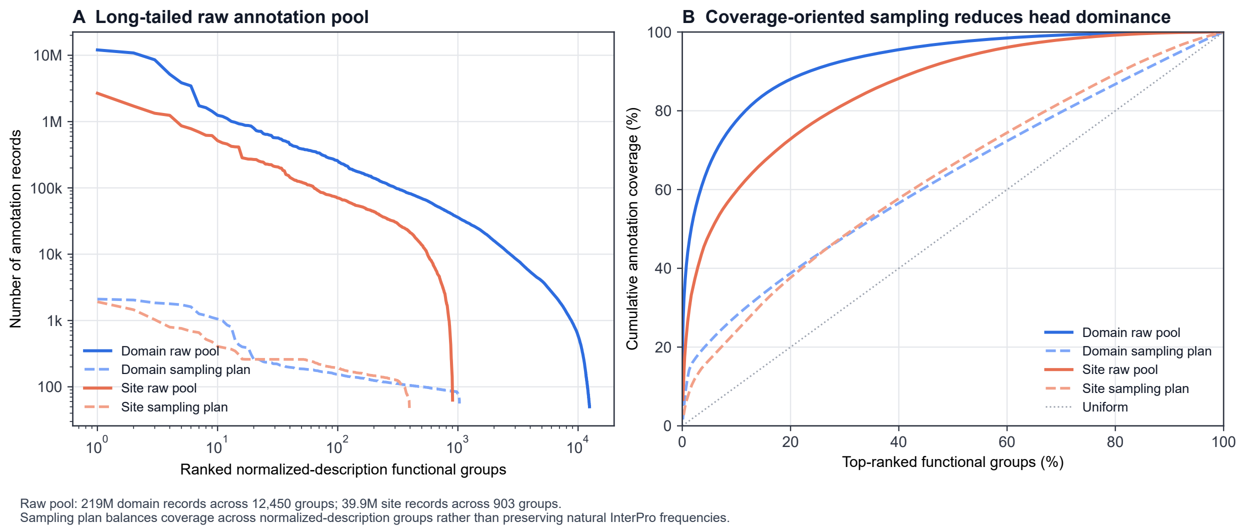


Figure S1: **Long-tailed distribution of normalized-description functional groups and coverage-oriented sampling.** (A) Rank-frequency curves of raw InterPro-derived annotation counts across normalized-description functional groups. Groups are ranked by annotation count within each category. Both domain and functional-site annotations show strong head–tail imbalance, especially for domain annotations. Dashed curves show the records retained by the coverage-oriented sampling plan. (B) Cumulative annotation coverage by top-ranked functional groups. In the raw pool, the top 1% of domain groups cover 44.1% of domain annotations and the top 10% cover 77.5%; for functional-site groups, the top 10% cover 59.8% of functional-site annotations. The sampling plan reduces head-group dominance and distributes selected records more evenly across functional groups, rather than preserving the natural InterPro frequency distribution.

We organized InterPro identifiers into normalized-description functional groups before sampling. These groups were not produced by embedding-based clustering or an automated semantic similarity model. Instead, they were constructed from normalized functional descriptions and manually curated high-frequency description patterns. Specifically, functional descriptions were normalized by lower-casing, removing redundant modifiers, regularizing punctuation and harmonizing common expression variants. Recurring description patterns that clearly referred to the same broad functional category, such as variants of conserved-site, binding-site, active-site or domain-family descriptions, were then consolidated into the same normalized-description functional group.

For domain annotations, identifiers were sampled across normalized-description functional groups to reduce the dominance of highly frequent domain families while retaining sufficient support for training and evaluation. For functional-site annotations, sampling was additionally stratified by site subtype, including `Active_site`, `Binding_site`, `Conserved_site` and `PTM`, to maintain coverage of different functional-site categories. The normalized-description functional groups and subtype labels were used only during benchmark construction and were not provided to the model during training or evaluation.

This description-group-aware sampling procedure selected 1,200 domain InterPro identifiers and 450 functional-site InterPro identifiers. These identifiers define the functional identifier set used for candidate protein and region collection.

##### S1.3 Candidate protein and region selection

After description-based identifier sampling, we collected candidate proteins and annotated regions associated with the selected InterPro identifiers. For each selected domain or functional-site identifier, we retrieved full-length proteins containing at least one retained annotation record for that identifier. Candidate collection was performed on full-length protein sequences rather than isolated fragments, because the downstream task requires localizing a functional region within a full-length protein sequence.

Within each protein–InterPro pair, duplicated annotation hits and near-duplicate regions were merged using the deterministic preprocessing rules described in Section S1.1. This reduced redundant annotation records while keeping the original InterPro residue coordinates for the retained representative spans.

Because some selected InterPro identifiers still had substantially larger support than others, candidate selection applied a per-identifier protein quota to limit the dominance of highly enriched identifiers. For each selected identifier, we retained at most 300 candidate proteins for domain identifiers and at most 260 candidate proteins for functional-site identifiers. When an identifier had more available proteins than the quota, candidate proteins were ranked by the quality score of their best retained candidate region, with the protein identifier used as a deterministic tie-breaker. This prioritized proteins with higher-quality annotated regions while keeping the selection reproducible. For functional-site annotations, candidate selection also preserved subtype coverage across `Active_site`, `Binding_site`, `Conserved_site` and `PTM`. Exact duplicate full-length sequences were then collapsed using sequence hashes, while retaining the associated source protein identifiers and candidate regions.

These records were then converted into explicit protein–text–region examples after the per-pair retention step described in Section S1.4.

##### S1.4 Explicit protein–text–region example construction

After candidate protein and region selection, we converted the retained candidate annotations into explicit protein–text–region examples. For each retained annotation record, the functional description associated with its InterPro identifier was used as the text query, and the original InterPro residue coordinates of the retained representative span were used as the localization target. Thus, the final supervision targets correspond to the original InterPro annotation spans rather than padded context windows.

To reduce repeated supervision from redundant annotations, we applied a per-pair retention rule after duplicate and near-duplicate records had been merged. For each protein–InterPro pair, we retained at most one domain span and up to two functional-site spans. The domain cap prevents repeated domain copies with highly similar functional descriptions from over-representing a small number of architectures in the benchmark. In contrast, up to two functional-site spans were retained because separated functional sites can correspond to the same functional query and are useful for evaluating ranked span proposal recovery.

This construction produced a pre-split explicit positive pool of 474,580 protein–text–region examples, including 344,445 domain examples and 130,135 functional-site examples. These examples were used as the candidate pool for sequence-similarity-aware split construction and subsequent benchmark materialization. During training, each retained example was treated independently as a single-region supervised instance. No explicit multi-site group label or set-level supervision was provided to the model. Visible multi-site grouping was constructed only for evaluation after the final data split, as described in Section S1.6.

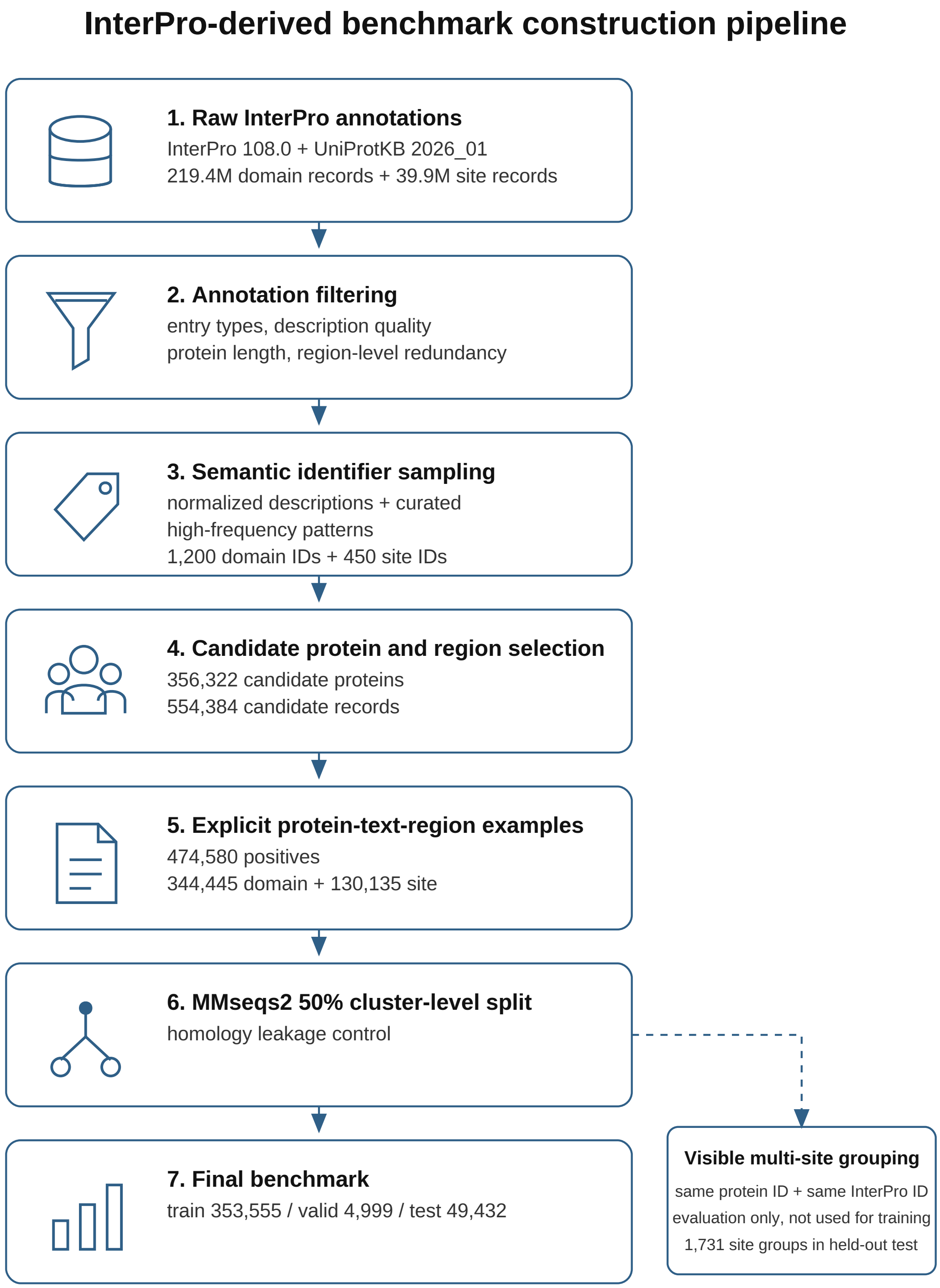


Figure S2: Overview of the InterPro-derived benchmark construction pipeline. InterPro annotations from InterPro 108.0 were linked to UniProtKB 2026_01 protein sequences and filtered by entry type, description quality, protein length and duplicate or near-duplicate region records. Functional identifiers were selected using description-group-aware sampling based on normalized descriptions and curated high-frequency description patterns. Candidate full-length proteins and functional-region records were then converted into explicit protein–text–region examples after per-pair retention, followed by MMseqs2 50% sequence-identity cluster-level splitting. Visible multi-site groups were constructed only for evaluation and were not used as training supervision.

Table S1: Final benchmark split statistics.

| Split | Samples | Proteins | Domain | Site | InterPro IDs |
|---|---|---|---|---|---|
| Train | 353,555 | 268,334 | 256,727 | 96,828 | 1,650 |
| Valid | 4,999 | 4,804 | 3,717 | 1,282 | 1,477 |
| Test | 49,432 | 34,219 | 36,547 | 12,885 | 1,477 |

##### S1.5 Sequence-similarity-aware data splitting

To reduce homology leakage between training and evaluation, we performed data splitting at the protein-cluster level rather than at the individual example level. This is necessary because multiple examples can be derived from the same protein, and the same InterPro identifier can appear in many homologous proteins. A random example-level split could therefore place identical or highly similar proteins in both training and test sets, leading to over-optimistic localization performance.

We clustered all candidate full-length protein sequences using MMseqs2 (Steinegger and Söding, 2017) with a 50% sequence identity threshold. Each resulting sequence-similarity cluster was assigned entirely to one split, so proteins from the same cluster could not appear in different splits. The split assignment was performed after explicit protein–text–region example construction, but the grouping constraint was defined by full-length protein sequence clusters.

During split construction, we balanced several factors, including the number of proteins, the number of explicit protein–text–region examples, the domain/site ratio and the distribution of functional-site subtypes. We also audited InterPro identifier support across splits so that validation and test metrics were not dominated by identifiers with very limited evaluation support. Additional identifiers were allowed only in the training split to increase training diversity; these train-only identifiers were excluded from validation and test metrics.

After protein-cluster-level split construction, split auditing and size control, 452,708 examples were retained in the full split materialization, including 353,555 training examples, 49,721 full-validation examples and 49,432 held-out test examples. The difference between the 474,580 pre-split explicit positive examples and the 452,708 full split examples comes from split construction, protein-cluster-level leakage control and training-size control. For efficient model selection, we then sampled a 4,999-example validation subset from the full-validation split. All checkpoint selection and hyperparameter choices were based only on this validation subset. The held-out test set containing 49,432 examples was used only after model selection for final benchmark reporting. The final released split statistics are reported in Section S1.6.

This cluster-level splitting strategy reduces direct sequence-similarity leakage by evaluating localization performance on proteins outside the training sequence-similarity clusters, while still maintaining sufficient functional-identifier and functional-site subtype coverage for reliable benchmark comparison.

##### S1.6 Final benchmark statistics and visible multi-site groups

The final released benchmark contains 407,986 explicit protein–text–region examples, including 353,555 training examples, 4,999 validation examples and 49,432 held-out test examples. This count corresponds to the released train/validation/test benchmark used for model selection and final evaluation. It differs from the 474,580-example pre-split explicit positive pool because the full split materialization retained 452,708 examples after split construction, leakage control and size control, and the 49,721-example full-validation split was further downsampled to the 4,999 validation examples used for checkpoint selection. The validation subset was used for checkpoint

Table S2: Protein–InterPro group composition in the held-out test set after final per-pair retention. Groups are defined by unique (protein_id, interpro_id) pairs.

| Region type | Records | Groups | Single-span groups | Multi-span groups |
|---|---|---|---|---|
| Domain | 36,547 | 36,547 | 36,547 | 0 |
| Site | 12,885 | 11,154 | 9,423 | 1,731 |
| Total | 49,432 | 47,701 | 45,970 | 1,731 |

selection and hyperparameter tuning, whereas the held-out test set was used only after model selection for final benchmark comparison. The split statistics are summarized in Table S1.

In addition to single-region evaluation, we constructed visible multi-site groups for evaluating ranked span proposals. A visible multi-site group is defined as a set of retained annotation records that share the same protein identifier and InterPro identifier and contain multiple retained target spans in the benchmark. These groups were constructed only after the final data split and were used only for evaluation. During training, each protein–text–region example was treated independently, and no multi-site group label or set-level supervision was provided to the model.

Table S2 reports the held-out test-set composition at the protein–InterPro group level. Groups are defined by unique (protein_id, interpro_id) pairs. The held-out test set contains 47,701 protein–InterPro groups, including 45,970 single-span groups and 1,731 multi-span groups. All multi-span groups come from functional-site annotations. Domain records have a one-to-one correspondence with protein–InterPro groups because the final retention rule keeps at most one domain span for each protein–InterPro pair. Functional-site annotations retain up to two spans per protein–InterPro pair, resulting in 1,731 visible multi-site groups, which account for 15.52% of functional-site groups and 3.63% of all held-out test groups.

The retention policy was designed to control redundancy and define a consistent evaluation target. Repeated domain copies within the same protein–InterPro pair often correspond to duplicated domain architectures and may over-represent a small number of repeated domain arrangements if all copies are retained. Therefore, the benchmark retains at most one domain span per protein–InterPro pair. In contrast, separated functional-site annotations can correspond to the same functional query and provide a setting for evaluating whether a model can recover multiple query-relevant regions, so up to two functional-site spans are retained.

We use the term "visible" because evaluation considers only the annotated regions retained in the benchmark. Potential functional regions that are not annotated in InterPro or not retained by the benchmark construction procedure are not treated as negatives. Therefore, the visible multi-site metrics evaluate recovery of benchmark-visible repeated functional-site annotations, rather than exhaustive discovery of all possible functional sites in a protein.

#### S2. Window-based baseline adaptation

##### S2.1 Baseline model families

We evaluated representative pretrained protein and protein–text models under a unified window-based grounding protocol. The evaluated baselines include Raw ESM2-650M (Lin et al., 2023), ProtST (Xu et al., 2023), ProtBERT, ProtT5 (Elnaggar et al., 2021), ProtCLIP (Zhou et al., 2025) and ProTrek (Su et al., 2025). These baselines cover three common categories of pretrained representations: protein-only language models, protein–text pretrained models and broader protein-related multimodal models.

Raw ESM2, ProtBERT and ProtT5 were used as protein-only representation baselines. Since these models do not natively include a text encoder, we paired each protein encoder with PubMed-BERT as the text tower and trained lightweight region–text alignment modules on the benchmark training set. This setting tests whether strong protein sequence representations can be adapted to text-conditioned functional region localization through region-level contrastive learning.

ProtST and ProtCLIP were used as protein–text pretrained baselines. These models already contain cross-modal pretraining signals between protein sequences and textual functional descriptions. We therefore adapted them using the same region–text contrastive protocol as the other baselines, so that their pretrained protein–text alignment could be evaluated under the proposed localization benchmark.

ProTrek was included as a sequence–structure–text multimodal protein baseline because it incorporates cross-modal pretraining among protein sequence, structure and natural-language functional descriptions.

All window-based baselines used the same training split, validation-based checkpoint selection and held-out test evaluation. Thus, their localization interface was restricted to window ranking rather than residue-level boundary prediction. Their predictions were obtained only by ranking candidate sequence windows according to region–text similarity.

##### S2.2 Region–text contrastive adaptation

We adapted all window-based baselines using the same region–text contrastive learning protocol. For each training example, the input consists of a full-length protein sequence, a functional text description and the annotated target span. The protein encoder first produces residue-level representations for the input protein sequence. The representations within the annotated span are then aggregated into a single region embedding using a lightweight trainable attention-pooling module. In parallel, the text encoder maps the corresponding functional description into a text embedding.

The region embedding and text embedding are projected into a shared representation space and L2-normalized before contrastive training. We used a CLIP-style symmetric contrastive objective (Zhou et al., 2025), where matched region–text pairs are treated as positives and mismatched region–text pairs within the same mini-batch are treated as negatives. A learnable temperature parameter is used to scale the similarity logits.

To reduce false-negative effects (Chuang et al., 2020), region–text pairs with the same normalized functional description were masked from the negative set. This is necessary because different proteins or regions may share the same functional description, and treating such pairs as negatives would incorrectly penalize query-equivalent region–text matches. This masking was applied uniformly to all window-based baselines.

Only lightweight task-adaptation components were optimized during baseline adaptation, including projection layers, the region attention-pooling module and the temperature parameter. The pretrained protein and text encoders were kept frozen. This design preserves the original pretrained representations while providing a unified lightweight mechanism for adapting different baselines to region-level protein–text matching.

After contrastive adaptation, each baseline can score an arbitrary candidate sequence window against the functional text query. These scores are used during sliding-window inference to rank candidate spans, as described in Section S2.3.

##### S2.3 Sliding-window inference

At inference time, the adapted baselines do not have access to ground-truth region coordinates and do not natively output text-conditioned residue boundaries. We therefore evaluated all window-

Table S3: Coverage of annotated span lengths by the maximum sliding-window size in the held-out test set.

| Region type | Number of spans | Spans $\leq$ 390 aa | Coverage |
|---|---|---|---|
| All | 49,432 | 49,244 | 99.62% |
| Domain | 36,547 | 36,357 | 99.48% |
| Site | 12,885 | 12,885 | 100.00% |

Table S4: Length distribution of site annotations in the held-out test set.

| Length threshold | Number of site spans | Fraction of site spans |
|---|---|---|
| $\leq$ 15 aa | 6,696 | 51.97% |
| $\leq$ 30 aa | 12,254 | 95.10% |
| $\leq$ 390 aa | 12,885 | 100.00% |

based baselines using the same multi-scale sliding-window retrieval protocol. Given a full-length protein sequence and a functional text query, candidate sequence windows were generated along the protein sequence and ranked by the learned region–text similarity score.

All window-based baselines used the same fixed multi-scale window set:

$$W = \{15, 30, 45, \dots, 390\}.$$

For each window size $w$, candidate windows were generated using a proportional stride:

$$\text{stride}(w) = \max(1, \lfloor 0.2w \rfloor).$$

Window sizes larger than the protein sequence length were skipped.

For each candidate window, residue representations inside the window were aggregated using the same attention-pooling module used during contrastive adaptation. The resulting window embedding and the text embedding were projected into the shared representation space and L2-normalized. The scaled cosine similarity between the two embeddings was used as the window score.

For each protein–text query pair, candidate windows were sorted in descending order of similarity. The highest-scoring window was used as the top-1 prediction for IoU@1 and boundary-based metrics. The ranked window list was used for Best@10 and visible multi-site metrics, with metric-specific cutoffs defined in Section S4.4. All predicted windows were represented as 0-based half-open spans and evaluated using the same span-level evaluator as ProLoc outputs.

The window range was chosen to cover both short functional sites and longer domain regions. As shown in Table S3, 99.62% of all annotated spans, 99.48% of domain spans and 100% of functional-site spans in the held-out test set have lengths no greater than 390 residues. The lower end of the window set was included to support short functional-site localization. As shown in Table S4, 51.97% of site spans are no longer than 15 residues, and 95.10% are no longer than 30 residues.

This protocol does not introduce a residue-level classifier, boundary regressor or anchor-free proposal head for the baselines. Therefore, under this fixed candidate-window protocol, baseline localization performance mainly reflects region–text alignment over a shared window grid rather than task-specific learned localization modules.

##### S2.4 Baseline checkpoint selection and hyperparameters

All window-based baselines were trained on the benchmark training set and selected using the validation subset. The held-out test set was used only after model selection for final benchmark evaluation. Each baseline was trained for at most 10 epochs and was periodically evaluated using the same sliding-window validation protocol described in Section S2.3. The final checkpoint for each baseline was selected according to validation Best@10 rather than the last training epoch, because Best@10 reflects the quality of the ranked window set used by the window-based grounding protocol.

Unless limited by GPU memory, baseline adaptation used 4 GPUs with a per-GPU batch size of 64, resulting in an effective global contrastive batch size of 256 after differentiable all-gather. For larger encoders, the per-GPU batch size was reduced when necessary due to memory constraints, while keeping the same contrastive objective, validation protocol and test evaluator.

All baselines were optimized using AdamW (Loshchilov and Hutter, 2017) with a learning rate of $5 \times 10^{-4}$, weight decay of 0.01 and gradient clipping for training stability. During adaptation, pretrained protein encoders and text encoders were kept frozen. Only lightweight task-adaptation components were optimized, including projection heads, the region attention-pooling module and the learnable temperature parameter.

For proteins longer than the maximum input length supported by the corresponding encoder, the training input was cropped around the annotated functional region and the target coordinates were shifted accordingly. This ensured that the complete ground-truth region remained visible to the model during contrastive adaptation. This cropping was used only during contrastive adaptation; during evaluation, no ground-truth coordinates were used, and all baselines followed the full sliding-window inference protocol and unified span-level evaluator.

#### S3. ProLoc training and implementation details

##### S3.1 Model configuration

The main ProLoc model uses raw ESM2-650M (Lin et al., 2023), i.e. the pretrained ESM2 sequence encoder without protein–text pretraining, as the protein encoder and PubMedBERT (Gu et al., 2021) as the text encoder. For each protein–text pair, the protein encoder provides residue-level sequence representations, while the text encoder provides both a pooled text representation and token-level text states. The pooled text representation is used for global text conditioning, and the token-level text states are used for residue–text interaction in the shared localization backbone.

The model contains a shared text-guided residue backbone followed by two output heads. The direct localization head predicts dense residue-level in-span logits. The anchor-free proposal head (Tian et al., 2019) predicts start, end and length signals for ranked span proposal generation. Both heads operate on the same shared residue representation, but they provide different output interfaces: dense residue-map localization and structured span proposal generation.

The direct-only ProLoc variant removes the anchor-free branch and keeps only the shared backbone and direct localization head. It is used as an ablation variant for isolating the effect of the anchor-free branch and its auxiliary supervision.

We also evaluate Raw-FiLM as a supervised dense control baseline. Raw-FiLM uses the same frozen raw ESM2-650M and PubMedBERT encoders as ProLoc, applies only pooled-text FiLM conditioning to residue representations, and is trained with the same direct localization loss and direct decoding protocol, without token-level residue-to-text cross-attention or anchor-free proposal learning.

##### S3.2 Training schedule

We trained two ProLoc variants separately. Direct-only ProLoc was trained with the shared text-guided residue backbone and the direct localization head only. It was optimized using only the direct localization objective and serves as an ablation for isolating dense residue-level supervision without the anchor-free branch.

Full ProLoc was trained as a separate model with both the direct localization head and the anchor-free proposal head. It was optimized using the direct localization objective together with anchor-free auxiliary objectives for start, end and length prediction. The full model was not obtained by continuing training from the Direct-only ProLoc checkpoint; it was trained independently with the complete multi-head objective.

During evaluation, the full ProLoc checkpoint produces two output forms. ProLoc-Direct denotes the output obtained by decoding the dense residue-level score map from the direct head, whereas ProLoc-AF denotes the ranked proposal output obtained by decoding the anchor-free start/end/length predictions together with the direct-head in-span confidence used for proposal scoring. Thus, ProLoc-Direct and ProLoc-AF are two decoding outputs from the same full model checkpoint, while Direct-only ProLoc is a separately trained ablation model.

All ProLoc variants were trained on the benchmark training set and selected using the validation subset. Checkpoints were evaluated periodically on the validation subset using the unified span-level evaluator. The held-out test set was used only for final benchmark comparison after model selection.

##### S3.3 Training objectives and loss weights

For each training example, the annotated functional region is represented as a 0-based half-open span $[s, e)$. The span is converted into a residue-level binary mask $y \in \{0,1\}^L$, where residues inside the target span are treated as positives and all other valid residues are treated as annotation-relative negatives for the retained training example. Padding positions are excluded from all loss computations.

The direct localization head is supervised by a weighted binary cross-entropy loss and a soft-IoU loss (Rahman and Wang, 2016). Given dense in-span logits $u = (u_1, \dots, u_L)$, we compute residue probabilities $p_i = \sigma(u_i)$. The binary cross-entropy term is computed over valid residues. To address residue-level class imbalance, the positive weight is set to $N_{\text{neg}}/N_{\text{pos}}$ for each training example and capped at 200. The soft-IoU loss is computed on the predicted probabilities and the binary target mask:

$$\mathcal{L}_{\text{softIoU}} = 1 - \frac{\sum_i p_i y_i}{\sum_i p_i + \sum_i y_i - \sum_i p_i y_i + \epsilon},$$

where the summations are over valid residues. The direct localization loss is

$$\mathcal{L}_{\text{dir}} = \mathcal{L}_{\text{BCE}}(u, y) + \lambda_{\text{IoU}} \mathcal{L}_{\text{softIoU}}(p, y).$$

where $\lambda_{\text{IoU}} = 0.5$.

The anchor-free branch uses three auxiliary losses for start, end and length prediction. For the start and end heads, we construct Gaussian-shaped soft boundary targets. The start target is centered at the ground-truth start position $s$, and the end target is centered at $e - 1$, corresponding to the last residue in the half-open target span. The Gaussian width is set to $\sigma = 2.0$. The start and end heads are trained with focal binary cross-entropy:

$$\mathcal{L}_{\text{start}} = \mathcal{L}_{\text{focal}}(a^{\text{start}}, g_s), \qquad \mathcal{L}_{\text{end}} = \mathcal{L}_{\text{focal}}(a^{\text{end}}, g_{e-1}),$$

Table S5: Training objectives and loss hyperparameters for ProLoc.

| Component | Setting | Value |
|---|---|---|
| Direct BCE | Positive weight | $N_{\text{neg}}/N_{\text{pos}}$, capped at 200 |
| Direct soft-IoU | Weight $\lambda_{\text{IoU}}$ | 0.5 |
| Start target | Center | $s$ |
| End target | Center | $e - 1$ |
| Boundary targets | Gaussian width $\sigma$ | 2.0 |
| Start/end focal loss | $\gamma$ | 2.0 |
| Start/end focal loss | $\alpha$ | 0.25 |
| Start loss | Weight $\lambda_s$ | 0.5 |
| End loss | Weight $\lambda_e$ | 0.5 |
| Length target | Log-length | $\log(\max(e - s, 1))$ |
| Length loss | Supervision position | ground-truth start $s$ only |
| Length loss | Weight $\lambda_l$ | 0.1 |

where $g_s$ and $g_{e-1}$ are the start and end soft boundary targets. The focal loss uses $\gamma = 2.0$ and $\alpha = 0.25$.

The length head is supervised only at the ground-truth start position. The target log-length is defined as

$$\ell^{\text{gt}} = \log(\max(e - s, 1)).$$

The length loss is an L1 loss at the start position:

$$\mathcal{L}_{\text{len}} = |\ell_s - \ell^{\text{gt}}|.$$

We do not apply length supervision to all residues because the predicted span length has a well-defined interpretation only when the residue is treated as a valid start position.

The full ProLoc objective combines the direct localization loss and the anchor-free auxiliary losses:

$$\mathcal{L} = \mathcal{L}_{\text{dir}} + \lambda_s \mathcal{L}_{\text{start}} + \lambda_e \mathcal{L}_{\text{end}} + \lambda_l \mathcal{L}_{\text{len}}.$$

We set $\lambda_s = 0.5$, $\lambda_e = 0.5$ and $\lambda_l = 0.1$. Direct-only ProLoc is trained only with $\mathcal{L}_{\text{dir}}$, whereas full ProLoc is trained with the complete objective above. During training, each protein–text–region record is treated as an independent single-region supervision example; no explicit multi-site group labels or set-level objectives are used. The main loss hyperparameters are summarized in Table S5.

##### S3.4 Optimization and implementation details

Both Direct-only ProLoc and full ProLoc were trained for 10 epochs on the benchmark training set. The protein encoder and text encoder were kept frozen during training. Only the task-specific localization modules were optimized, including the text-guided fusion layers, the direct localization head and, for full ProLoc, the anchor-free proposal branch.

Epoch-level checkpoints were evaluated on the validation subset using the unified span-level evaluator. The held-out test set was used only after checkpoint selection for final benchmark reporting. The final held-out test evaluation was performed using 4-GPU distributed inference. Checkpoint selection was based on validation Best@10.

The main optimization and implementation settings are summarized in Table S6.

Table S6: Optimization and implementation settings for ProLoc.

| Setting | Value |
|---|---|
| Training epochs | 10 |
| Optimizer | AdamW |
| Learning rate | $5 \times 10^{-4}$ |
| Weight decay | 0.01 |
| Gradient clipping | max norm 1.0 |
| Protein encoder | frozen raw ESM2-650M |
| Text encoder | frozen PubMedBERT |
| Training batch size | 64 |
| Validation / test batch size | 16 |
| Checkpoint selection | validation subset |
| Final test inference | 4-GPU distributed inference |
| Framework | PyTorch / ProtST codebase |
| Python | 3.9.25 |
| PyTorch | 2.0.0 |
| Transformers | 4.30.2 |
| NumPy | 1.24.4 |
| scikit-learn | 1.6.1 |

#### S4. Unified decoding and evaluation protocol

##### S4.1 Output normalization into top-$K$ spans

Different methods produce different native outputs. Window-based baselines rank predefined sequence windows by region–text similarity. The direct head of ProLoc produces dense residue-level in-span scores, whereas the anchor-free head produces start, end and length signals for proposal generation. To ensure a consistent comparison across these heterogeneous output formats, we normalize all method outputs into the same ranked span representation before computing evaluation metrics.

For each protein–text query pair, the normalized prediction is represented as

$$\hat{\mathcal{S}} = \{(\hat{s}_k, \hat{e}_k, \hat{c}_k)\}_{k=1}^{K},$$

where $(\hat{s}_k, \hat{e}_k)$ denotes the $k$-th predicted 0-based half-open span and $\hat{c}_k$ denotes its confidence score. Spans are sorted in descending order of $\hat{c}_k$. If a method produces fewer than $K$ valid spans, all available spans are retained. Unless otherwise specified, we use $K = 10$ for top-$K$ evaluation.

For window-based baselines, the ranked spans are the candidate sequence windows sorted by region–text similarity. For the direct head, ranked spans are obtained by decoding connected components from the dense residue-level score map. For the anchor-free head, ranked spans are generated from predicted boundary and length signals, followed by proposal scoring and non-maximum suppression. The method-specific decoding procedures are described in Sections S4.2 and S4.3.

This normalization step decouples model-specific prediction mechanisms from metric computation. Single-region metrics, such as IoU@1 and B-MAE, use the highest-ranked span $(\hat{s}_1, \hat{e}_1)$. Top-$K$ and visible multi-site metrics use the ranked span list with the metric-specific cutoffs defined in Section S4.4. Therefore, all baselines and ProLoc variants are evaluated under the same span-level protocol despite having different native output forms.

##### S4.2 Direct-head decoding

The direct head outputs dense residue-level logits for each protein–text query pair:

$$u = (u_0, u_1, \dots, u_{L-1}).$$

We first convert the logits into residue-level probabilities using a sigmoid function:

$$p_i = \sigma(u_i), \qquad i = 0, \dots, L-1.$$

Before thresholding, the probability map is smoothed using a fixed centered one-dimensional moving-average window. In all experiments, the smoothing window size is set to 9 residues. Specifically, for each residue position $i$, we compute

$$\bar{p}_i = \frac{1}{|\mathcal{N}_i|} \sum_{j \in \mathcal{N}_i} p_j, \qquad \mathcal{N}_i = \{j : 0 \le j < L,\ |j - i| \le 4\}.$$

The smoothed probability map is denoted as $\bar{p}$. This fixed smoothing step reduces isolated noisy activations and stabilizes connected-component extraction.

We then apply a fixed threshold 0.5 to $\bar{p}$. Consecutive residues with $\bar{p}_i \ge 0.5$ are grouped into connected components. Each connected component is converted into a predicted 0-based half-open span $(\hat{s}, \hat{e})$. For each decoded span, the confidence score is computed as the mean smoothed probability within the component:

$$\hat{c}(\hat{s}, \hat{e}) = \frac{1}{\hat{e} - \hat{s}} \sum_{i=\hat{s}}^{\hat{e}-1} \bar{p}_i.$$

Decoded spans are sorted in descending order of $\hat{c}$, producing the ranked span list defined in Section S4.1.

If no residue passes the threshold, we use the highest-scoring residue in $\bar{p}$ as a fallback one-residue span, with ties broken by the smallest residue index. This guarantees that every protein–text query pair has a valid top-1 prediction. Direct-head decoding uses only the predicted score map and fixed decoding hyperparameters; it does not use ground-truth coordinates during inference.

##### S4.3 Anchor-free proposal decoding

The anchor-free head predicts boundary and span-size signals for each valid residue. For residue $i = 0, \dots, L-1$, the model outputs a start logit, an end logit and a predicted log-length:

$$a_i^{\text{start}}, \qquad a_i^{\text{end}}, \qquad \ell_i.$$

The start and end logits are converted into probabilities:

$$p_i^{\text{start}} = \sigma(a_i^{\text{start}}), \qquad p_i^{\text{end}} = \sigma(a_i^{\text{end}}).$$

Padding positions are excluded by the valid residue mask throughout proposal generation.

Candidate starts are selected from residues with $p_i^{\text{start}} \ge 0.3$. If more than 64 residues satisfy this threshold, only the top 64 residues ranked by $p_i^{\text{start}}$ are retained. If no residue satisfies the threshold, we fall back to the top 64 start positions over the full valid sequence. This fallback prevents empty predictions for difficult protein–text pairs and ensures that the anchor-free decoder can always generate candidate proposals.

We use two complementary proposal-generation modes. In the length-based mode, a candidate span is generated from a start position $s$ and its predicted log-length. The predicted log-length is first clipped to the range $[\log 2, \log 500]$, and the span length is computed as

$$m_s = \mathrm{round}(\exp(\ell_s)).$$

The resulting length is further clipped to the range $[2, 500]$. The length-based candidate span is then constructed as

$$[\hat{s}, \hat{e}) = [s, \min(s + m_s, L)).$$

In the end-pair mode, each candidate start $s$ is paired with high-confidence inclusive end residues. For each start $s$, candidate end residues are searched within the valid length range

$$r \in \{s + 2 - 1, \ldots, \min(s + 500, L) - 1\}.$$

End residues with $p_r^{\mathrm{end}} \geq 0.3$ are selected, and at most the top 8 end residues ranked by $p_r^{\mathrm{end}}$ are retained for each start. If no end residue in the allowed range satisfies the threshold, we fall back to the top 8 end residues within that range. Each start–end pair $(s, r)$ is converted into the 0-based half-open span

$$[s, r + 1).$$

The final candidate pool is the union of length-based proposals and end-pair proposals.

Each candidate span is assigned a confidence score using boundary evidence and dense in-span evidence from the direct head. Let $p_i^{\mathrm{dir}} = \sigma(u_i)$ denote the direct-head residue-level in-span probability. For a candidate span $[s, e)$, the average direct in-span confidence is

$$\bar{p}_{\mathrm{dir}}(s, e) = \frac{1}{e - s} \sum_{i=s}^{e-1} p_i^{\mathrm{dir}}.$$

For length-based proposals, the confidence score is

$$\hat{c}_{\mathrm{len}}(s, e) = p_s^{\mathrm{start}}\, \bar{p}_{\mathrm{dir}}(s, e).$$

For end-pair proposals, the confidence score is

$$\hat{c}_{\mathrm{pair}}(s, e) = p_s^{\mathrm{start}}\, p_{e-1}^{\mathrm{end}}\, \bar{p}_{\mathrm{dir}}(s, e).$$

Candidate spans from both modes are sorted by their confidence scores. If the merged candidate pool contains more than 256 spans, only the top 256 candidates are retained before non-maximum suppression.

To remove redundant proposals, we apply greedy non-maximum suppression based on span IoU. Unless otherwise specified, the NMS IoU threshold is set to 0.3: after sorting candidates by confidence score, any subsequent candidate with span IoU greater than 0.3 with a previously retained proposal is suppressed. The decoder then keeps the top 10 non-redundant spans as the anchor-free ranked proposal output. These spans are evaluated using the same ranked span protocol defined in Section S4.1.

Anchor-free decoding uses only predicted start, end, length and direct in-span scores together with fixed decoding hyperparameters. It does not use ground-truth coordinates during inference. Compared with direct-head decoding, this procedure explicitly models boundary and span-size evidence and provides structured top-$K$ proposals for visible multi-site evaluation.

##### S4.4 Evaluation metrics

All methods are evaluated using the normalized ranked span list defined in Section S4.1. For each protein–text query pair, the model outputs

$$\hat{\mathcal{S}} = \left[(\hat{s}_k, \hat{e}_k, \hat{c}_k)\right]_{k=1}^{K}$$

where predicted spans are sorted by confidence in descending order. We denote the $k$-th predicted span as $\hat{S}_k = [\hat{s}_k, \hat{e}_k)$. Unless otherwise specified, top-$K$ metrics use $K = 10$.

For single-region localization, each evaluation example has one ground-truth span $S = [s, e)$. The primary metric is the intersection-over-union between the top-ranked prediction and the ground truth:

$$\text{IoU@1} = \frac{|\hat{S}_1 \cap S|}{|\hat{S}_1 \cup S|}.$$

We also report Domain IoU@1 and Functional-Site IoU@1 by computing the same metric separately on domain and site examples.

To measure boundary accuracy, we report boundary mean absolute error:

$$\text{B-MAE} = \frac{|\hat{s}_1 - s| + |\hat{e}_1 - e|}{2}.$$

B-MAE is computed using the top-ranked predicted span $\hat{S}_1$, and lower values indicate more accurate boundary localization.

For top-$K$ localization quality, we report Best@10:

$$\text{Best@10} = \max_{1 \le k \le 10} \text{IoU}(\hat{S}_k, S).$$

Best@10 measures whether the ground-truth region appears among the top-ranked candidate spans even if it is not ranked first. Domain Best@10 and Functional-Site Best@10 are computed analogously on the corresponding subsets.

For visible multi-site evaluation, we group retained test records by the same protein identifier and InterPro identifier. Let $\mathcal{G}$ denote the set of visible multi-site groups. Each group $g \in \mathcal{G}$ contains $m_g$ benchmark-visible target spans:

$$\mathcal{T}_g = \{S^{(g,1)}, S^{(g,2)}, \ldots, S^{(g,m_g)}\}.$$

These groups are used only for evaluation and are not provided as training supervision. The term "visible" indicates that evaluation only considers annotated spans retained in the benchmark; unannotated or unretained potential functional regions are not treated as negatives.

Given the top-10 predicted spans for the corresponding protein–text query, a visible target span $S^{(g,j)}$ is considered recovered at IoU threshold $\tau$ if

$$\max_{1 \le k \le 10} \text{IoU}(\hat{S}_k, S^{(g,j)}) \ge \tau.$$

Visible multi-site recall is computed over all visible target spans:

$$\text{VM R@10 IoU}\tau = \frac{\sum_{g \in \mathcal{G}} \sum_{j=1}^{m_g} \mathbf{1}\left[\max_{1 \le k \le 10} \text{IoU}(\hat{S}_k, S^{(g,j)}) \ge \tau\right]}{\sum_{g \in \mathcal{G}} m_g}.$$

We report this metric at $\tau = 0.5$ and $\tau = 0.7$, denoted as VM R@10 IoU50 and VM R@10 IoU70, respectively.

We further report a group-level all-hit metric. A visible multi-site group is counted as successfully recovered only if every visible target span in the group is matched by at least one top-10 prediction with IoU at least 0.5:

$$\text{VM All-Hit@50} = \frac{1}{|\mathcal{G}|}\sum_{g\in\mathcal{G}} \mathbf{1}\left[\forall j \in \{1,\ldots,m_g\},\ \max_{1\le k\le 10} \text{IoU}(\hat{S}_k, S^{(g,j)}) \ge 0.5\right].$$

Finally, we report visible multi-site mean IoU:

$$\text{VM Mean IoU@10} = \frac{\sum_{g\in\mathcal{G}}\sum_{j=1}^{m_g} \max_{1\le k\le 10} \text{IoU}(\hat{S}_k, S^{(g,j)})}{\sum_{g\in\mathcal{G}} m_g}.$$

This metric measures the best overlap achieved by the top-10 predictions for each benchmark-visible target span without applying a binary IoU threshold.

Together, IoU@1 and B-MAE evaluate the highest-ranked localization result, Best@10 evaluates candidate-set quality, and the visible multi-site metrics evaluate whether ranked proposals can recover multiple benchmark-visible functional-site regions for the same protein–text query. Ground-truth coordinates are used only for metric computation and are never used during decoding.

### Supplementary Results

#### S5. Quantitative control analyses

To further examine whether the proposed model benefits from query-dependent localization rather than only dense residue-level supervision or fixed protein-level saliency, we report two additional control analyses. First, we evaluate Raw-FiLM, a supervised dense baseline using the same raw ESM2-650M and PubMedBERT encoders but only pooled-text FiLM conditioning. Second, we perform a matched wrong-query control, where each test protein is kept unchanged while the functional description is replaced by a type- and length-matched incorrect description. The evaluation target remains the original annotated span, so this control tests query dependence rather than absent-query detection.

Table S7: Full metrics of the Raw-FiLM supervised dense baseline. Raw-FiLM uses the same raw ESM2-650M and PubMedBERT encoders as ProLoc, but conditions residue representations only through pooled-text FiLM modulation, without token-level residue-to-text cross-attention or anchor-free proposal generation. Higher is better except for B-MAE.

| Method | Type | IoU@1 | Domain IoU@1 | Site IoU@1 | B-MAE ↓ | Best@10 | Domain Best@10 | Site Best@10 | VM R@10 IoU50 | VM R@10 IoU70 | VM All-Hit@50 | VM Mean IoU@10 |
|---|---|---|---|---|---|---|---|---|---|---|---|---|
| Raw-FiLM | Direct | 0.5568 | 0.6218 | 0.3725 | 99.59 | 0.6023 | 0.6606 | 0.4371 | 0.3645 | 0.1521 | 0.1810 | 0.3923 |

Table S8: Matched wrong-query control for ProLoc-Direct. For each test example, the protein sequence is kept unchanged, while the functional description is replaced by a type- and length-matched incorrect description. Metrics are still computed against the original annotated span. Higher is better except for B-MAE.

| Setting | IoU@1 | Domain IoU@1 | Site IoU@1 | B-MAE ↓ | Best@10 | Domain Best@10 | Site Best@10 | VM R@10 IoU50 | VM R@10 IoU70 | VM All-Hit@50 | VM Mean IoU@10 |
|---|---|---|---|---|---|---|---|---|---|---|---|
| True query | 0.7730 | 0.7938 | 0.7141 | 65.77 | 0.8142 | 0.8153 | 0.8163 | 0.8322 | 0.7602 | 0.7112 | 0.7597 |
| Matched wrong query | 0.1621 | 0.1089 | 0.3129 | 262.79 | 0.1930 | 0.1395 | 0.3448 | 0.2777 | 0.1900 | 0.1404 | 0.2779 |

#### S6. Qualitative and error analysis

##### S6.1 Qualitative localization examples

We provide representative qualitative localization examples in Fig. S3. These cases illustrate four representative settings from the held-out test set: long-domain localization, short functional-site localization, visible multi-site recovery and query-specific localization on the same protein sequence.

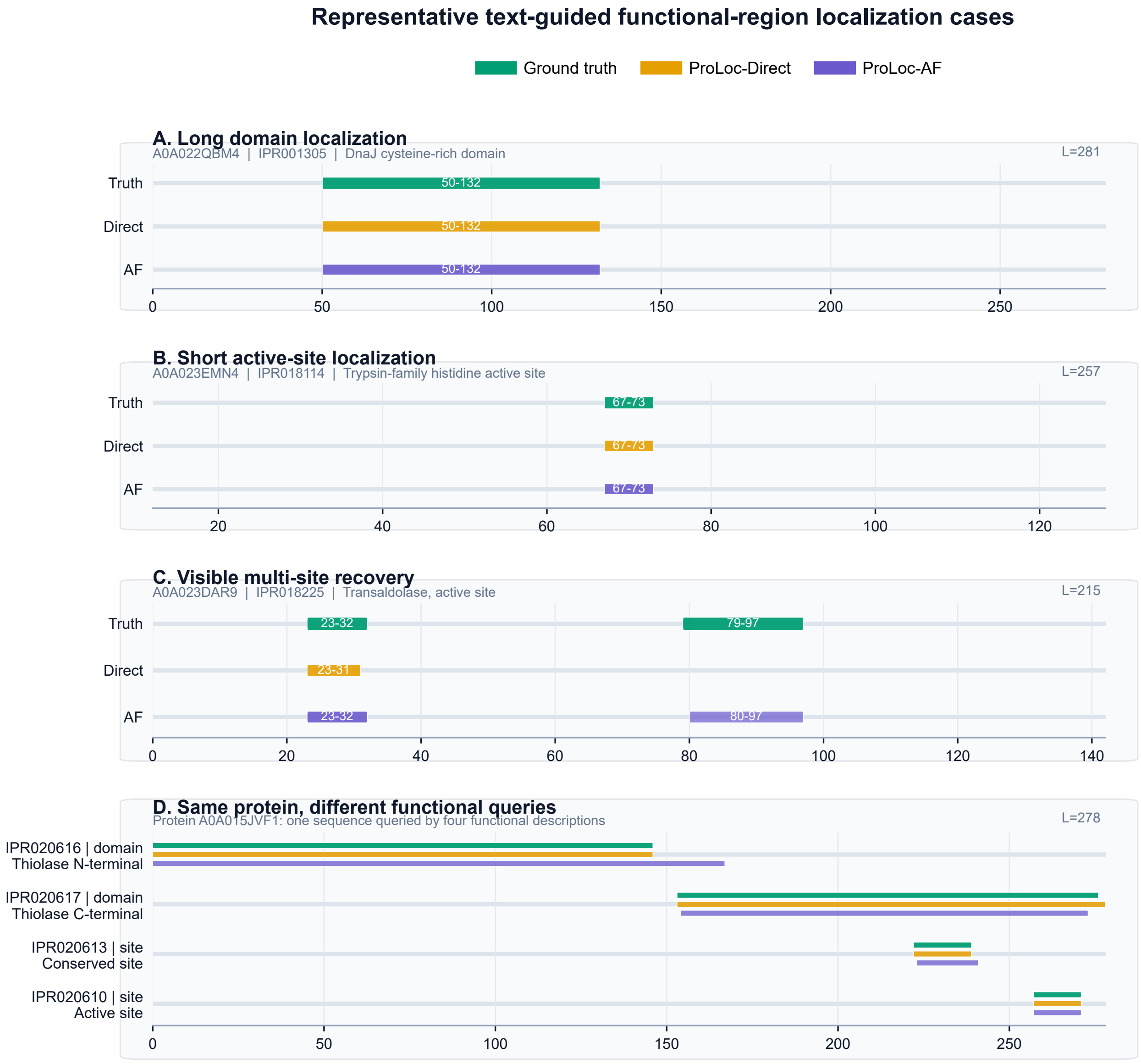


Figure S3: Representative text-guided functional-region localization examples. Each panel compares the ground-truth annotation with the ProLoc-Direct output and the ProLoc-AF proposal output. The examples cover long-domain localization, short functional-site localization, visible multi-site recovery and query-specific localization on the same protein sequence. Residue coordinates follow the 0-based half-open convention used throughout the benchmark.

For long functional regions, the qualitative example shows that ProLoc can recover domain-level spans accurately. In the DnaJ cysteine-rich domain example, both ProLoc-Direct and ProLoc-

AF align closely with the annotated domain region, illustrating that the text-conditioned residue representation can support localization of extended functional regions.

For short functional sites, ProLoc also preserves residue-level specificity. In the trypsin-family histidine active-site example, the annotated short site is recovered by both the direct output and the anchor-free proposal output. This example illustrates that the model can localize compact functional regions rather than only broad domain-scale regions.

The visible multi-site example shows the complementary role of the anchor-free proposal head. For the transaldolase active-site query, the ground truth contains two separated site regions in the same protein. ProLoc-Direct recovers the dominant connected region, whereas ProLoc-AF produces separated span proposals that cover both benchmark-visible sites. This behavior is consistent with the quantitative visible multi-site results, where the anchor-free output improves multi-site recall and all-hit rate.

Finally, the same-protein example demonstrates that ProLoc is query-conditioned rather than simply highlighting fixed regions of a protein sequence. For the same thiolase protein, different functional descriptions lead to different localized regions, including N-terminal and C-terminal thiolase domains as well as conserved and active sites. This supports the task formulation that the predicted functional region should change according to the input text query.

##### S6.2 Error analysis

We provide representative failure cases in Fig. S4. These examples illustrate representative remaining errors of ProLoc, including boundary shifts on short functional sites, incomplete recovery of separated regions and confusion between repeated or functionally related regions.

First, short functional sites are more sensitive to small boundary shifts than long domains. For a domain-scale region, a few-residue boundary offset usually has limited effect on IoU. In contrast, for a short active site or binding site, the same absolute offset can substantially reduce overlap. This helps explain why functional-site localization is more sensitive to boundary errors than domain localization and motivates reporting both IoU-based metrics and boundary error.

Second, the direct and anchor-free outputs show different failure modes. The direct head predicts a dense residue-level in-span map and usually provides the strongest top-1 localization. However, when the same functional query corresponds to multiple separated sites, thresholding the dense score map may emphasize the most confident connected component and miss weaker separated regions. The anchor-free output is designed for ranked multi-span proposal generation, but its top-1 prediction can be less precise when start, end or length estimates are slightly misaligned.

Third, repeated or homologous functional regions can introduce ambiguous localization cases. Some proteins contain repeated domains or multiple similar local motifs associated with the same or closely related functional description. In such cases, a prediction may highlight a plausible repeated or functionally related region that does not match the retained benchmark span. The benchmark evaluates overlap with the retained InterPro annotation coordinates, so such predictions are counted as errors if they do not sufficiently overlap the benchmark-visible target.

Finally, visible multi-site evaluation is limited to benchmark-retained annotations. The term "visible" indicates that only retained annotated spans are evaluated. Potential functional regions that are unannotated or removed during benchmark construction are not treated as negatives. Therefore, visible multi-site errors should be interpreted as failures to recover benchmark-visible repeated site annotations, rather than exhaustive failures to discover every possible functional site in a protein.

Overall, these observations highlight remaining challenges in compact site localization, boundary calibration, repeated-region disambiguation and multi-span proposal ranking. Future improvements may incorporate stronger boundary calibration, explicit set-level objectives for repeated sites and uncertainty-aware treatment of ambiguous or incomplete functional annotations.

#### References


M. Blum, H.-Y. Chang, S. Chuguransky, T. Grego, S. Kandasaamy, A. Mitchell, G. Nuka, T. Paysan-Lafosse, M. Qureshi, S. Raj, et al. The interpro protein families and domains database: 20 years on. *Nucleic acids research*, 49(D1):D344–D354, 2021.

C.-Y. Chuang, J. Robinson, Y.-C. Lin, A. Torralba, and S. Jegelka. Debiased contrastive learning. *Advances in neural information processing systems*, 33:8765–8775, 2020.

A. Elnaggar, M. Heinzinger, C. Dallago, G. Rehawi, Y. Wang, L. Jones, T. Gibbs, T. Feher, C. Angerer, M. Steinegger, et al. Prottrans: toward understanding the language of life through self-supervised learning. *IEEE transactions on pattern analysis and machine intelligence*, 44(10): 7112–7127, 2021.

Y. Gu, R. Tinn, H. Cheng, M. Lucas, N. Usuyama, X. Liu, T. Naumann, J. Gao, and H. Poon. Domain-specific language model pretraining for biomedical natural language processing. *ACM Transactions on Computing for Healthcare (HEALTH)*, 3(1):1–23, 2021.

Z. Lin, H. Akin, R. Rao, B. Hie, Z. Zhu, W. Lu, N. Smetanin, R. Verkuil, O. Kabeli, Y. Shmueli, et al. Evolutionary-scale prediction of atomic-level protein structure with a language model. *Science*, 379(6637):1123–1130, 2023.

I. Loshchilov and F. Hutter. Decoupled weight decay regularization. *arXiv preprint arXiv:1711.05101*, 2017.

J. Mistry, S. Chuguransky, L. Williams, M. Qureshi, G. A. Salazar, E. L. Sonnhammer, S. C. Tosatto, L. Paladin, S. Raj, L. J. Richardson, et al. Pfam: The protein families database in 2021. *Nucleic acids research*, 49(D1):D412–D419, 2021.

M. A. Rahman and Y. Wang. Optimizing intersection-over-union in deep neural networks for image segmentation. In *International symposium on visual computing*, pages 234–244. Springer, 2016.

M. Steinegger and J. Söding. Mmseqs2 enables sensitive protein sequence searching for the analysis of massive data sets. *Nature biotechnology*, 35(11):1026–1028, 2017.

J. Su, Y. He, S. You, S. Jiang, X. Zhou, X. Zhang, Y. Wang, X. Su, I. Tolstoy, X. Chang, et al. A trimodal protein language model enables advanced protein searches. *Nature Biotechnology*, pages 1–7, 2025.

Z. Tian, C. Shen, H. Chen, and T. He. Fcos: Fully convolutional one-stage object detection. In *Proceedings of the IEEE/CVF international conference on computer vision*, pages 9627–9636, 2019.

UniProt Consortium. Uniprot: the universal protein knowledgebase in 2025. *Nucleic Acids Research*, 53(D1):D609–D617, 2025. doi: 10.1093/nar/gkae1010.

M. Xu, X. Yuan, S. Miret, and J. Tang. Protst: Multi-modality learning of protein sequences and biomedical texts. In *International conference on machine learning*, pages 38749–38767. PMLR, 2023.

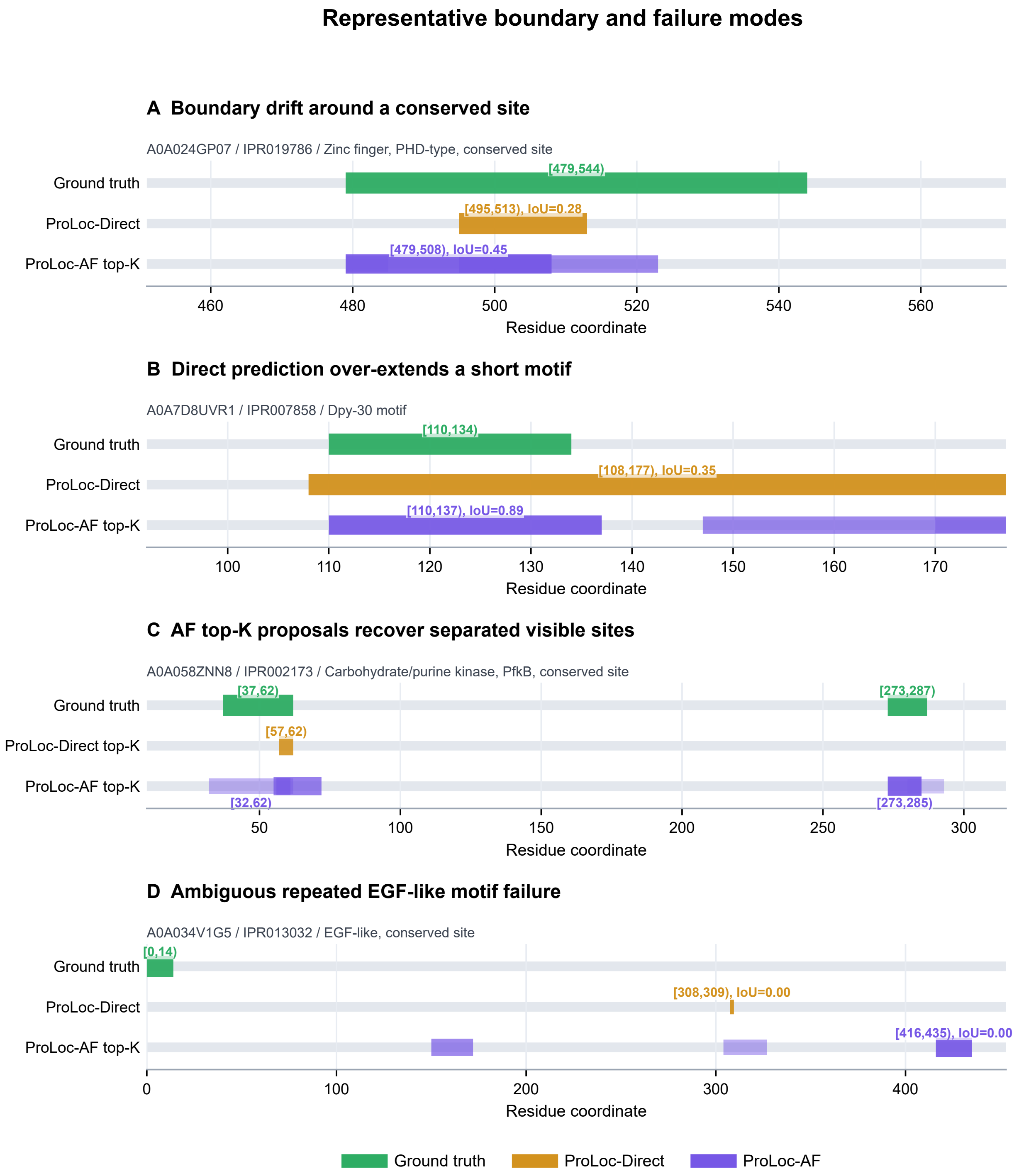


Figure S4: Representative failure cases of text-guided functional-region localization. The examples illustrate common error modes observed in the held-out test set, including boundary offsets for compact functional sites, partial recovery of separated visible functional sites and ambiguous localization among repeated or functionally related regions. Residue coordinates follow the 0-based half-open convention used throughout the benchmark.

H. Zhou, M. Yin, W. Wu, M. Li, K. Fu, J. Chen, J. Wu, and Z. Wang. Protclip: Function-informed protein multi-modal learning. In *Proceedings of the AAAI Conference on Artificial Intelligence*, volume 39, pages 22937–22945, 2025.
